# The phosphoinositide signature guides the final step of plant cytokinesis

**DOI:** 10.1101/2022.12.18.520917

**Authors:** Alexis Lebecq, Aurélie Fangain, Elsa Gascon, Camila Goldy, Katia Belcram, Martine Pastuglia, David Bouchez, Marie-Cécile Caillaud

## Abstract

Plant cytokinesis, which fundamentally differs from that in animals, involves de novo assembly of a plasma membrane precursor named the cell plate. How the transition from the cell plate to a plasma membrane occurs at the end of the plant cytokinesis remains poorly understood. Here, we describe with unprecedented spatiotemporal precision, the acquisition of plasma membrane identity upon cytokinesis through the lateral patterning of phosphatidylinositol 4,5-bisphosphate PI(4,5)P_2_ at the newly formed cell plate membrane. We show that during late cytokinesis, opposing polarity domains are formed along the cell plate. Exclusion of PI(4,5)P_2_ from the leading edge of the cell plate is controlled by SAC9, a putative phosphoinositide phosphatase. SAC9 colocalizes with MAP65-3, a key regulator of the cytokinesis, at the cell plate leading zone and regulates its function. In the *sac9-3* mutant, the polar distribution of PI(4,5)P_2_ at the cell plate is altered, leading to de-novo recruitment of the cytokinesis apparatus and to formation of an additional, ectopic cell plate insertion site. We proposed that PI(4,5)P_2_ acts as a polar cue to spatially separate the expansion and maturation domains of the forming cell plate during the final steps of cytokinesis.

**One Sentence Summary:** The phosphoinositide PI(4,5)P_2_ acts as an hallmark to guide the final step of plant cell division.

## INTRODUCTION

Plant cytokinesis differs from the inward cytokinesis observed in animals, and involves de novo building of a new cell plate between daughter cells, established in telophase by a plant-specific cytoskeletal array named the phragmoplast (*1*). This process requires directional vesicle trafficking toward the phragmoplast midzone, for the deposition of a transitory membrane structure named the cell plate (*2*). There, the cell plate expands toward the cell periphery as microtubules are disassembled in the inner phragmoplast region (lagging zone), while new overlapping antiparallel microtubules appear at the outer edge of the phragmoplast (leading zone, Fig. 1a). In this process, the most likely microtubule cross-linking factors are proteins belonging to the MAP65 family and members of the Kinesin-12 subfamily (*3, 4*). Upon attachment of the growing cell plate to the mother membrane, cell plate properties and composition undergo a number of changes (*5–7*), suggesting that the phragmoplast leading zone might act as a landmark to terminate cytokinesis (*8*). The nature of this cue remains elusive, and the molecular players regulating the transition from cell plate to plasma membrane identity are poorly understood.

**Fig.1.**
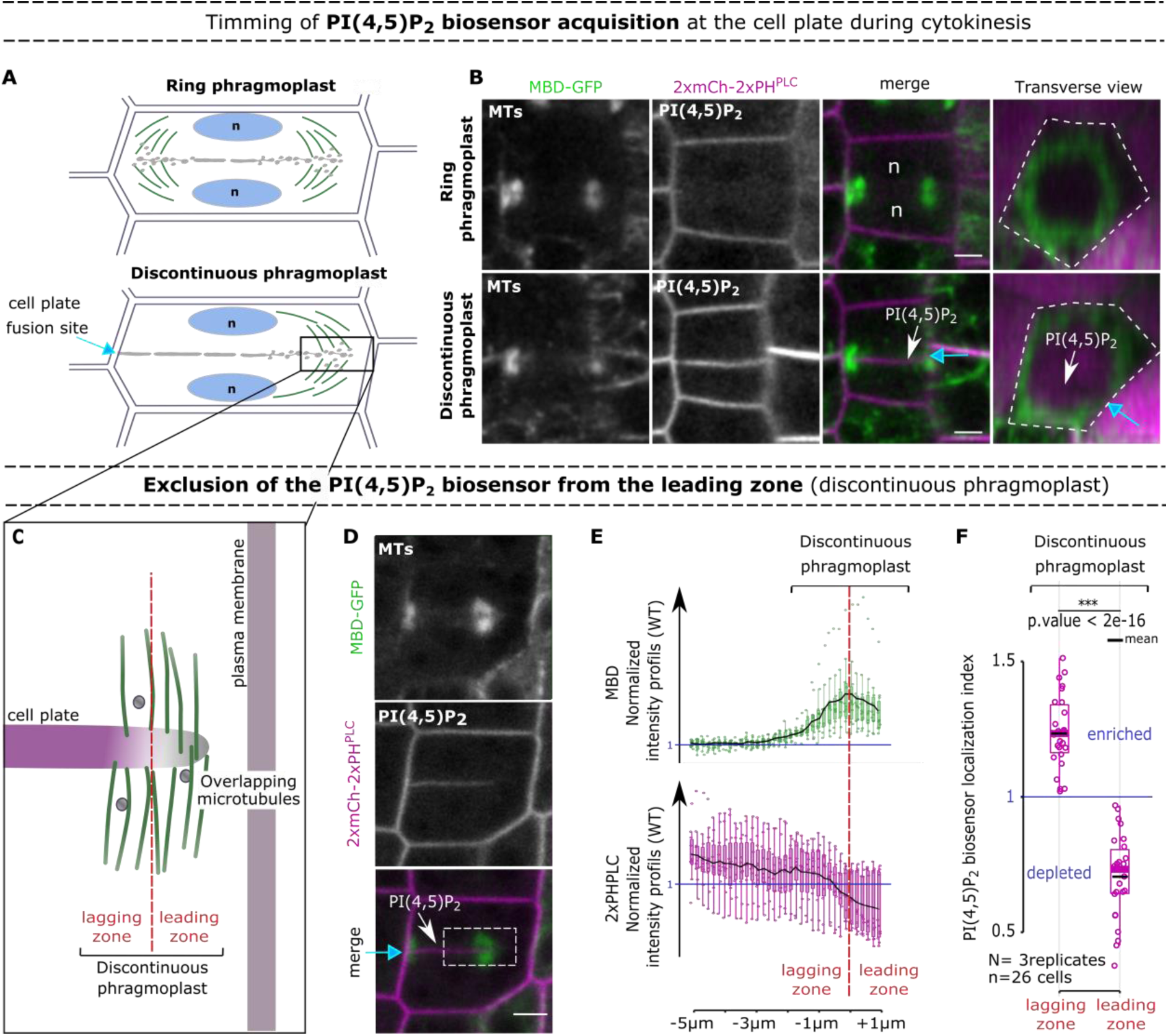
PI(4,5)P2 is recruited at the cell plate after its unilateral attachment. **a**, Representation of the unattached cell plate showing a ring phragmoplast (top) and the unilaterally attached cell plate corresponding to a discontinuous phragmoplast (bottom). **b**, Confocal images with a root tracking system (*38*) of 2xmCh-2xPH^PLC^ with MBD-GFP during the two steps represented in (a). Single longitudinal sections for each fluorescent channel and a transverse section of the merge channels is presented (0.7µm between optical sections). Dotted line, cell contour. **c**, Representation of the expanding edge of a discontinuous phragmoplast. **d**, Images of 2xmCh-2xPH^PLC^ and MBD-GFP at the unilaterally attached cell plate. **e**, Normalized intensity profiles at 6 µm along the discontinuous phragmoplast (ROI). **f**, PI(4,5)P_2_ biosensor localization index (normalized by the cell plate intensity) on the phragmoplast lagging/leading zones. In the plots, middle horizontal bars represent the median, while the bottom and top of each box represent the 25^th^ and 75^th^ percentiles, respectively. At most, the whiskers extend to 1.5 times the interquartile range, excluding data beyond. For range of value under 1,5 IQR, whiskers represent the range of maximum and minimum values. Results of the statistical analysis (shown in the supplementary table) are presented (N = number of replicates, n = number of cells). White arrow, PI(4,5)P_2_ biosensor appearance; Blue arrow, cell plate fusion site; red dotted line, separation leading and lagging zone; dotted lines, region of interest; n, nucleus; scale bars, 5 µm.

Recent evidence points toward a specific anionic lipid signature for the plant plasma membrane (*9*). This landmark is set up by enzymes involved in phosphoinositide metabolism, in particular phosphatases and kinases that locally convert phosphoinositide pools (*10*). At the plasma membrane, phosphatidylinositol 4,5-bisphosphate (PI(4,5)P_2_) is enriched, whereas PI(4,5)P_2_ is excluded from the endocytic pathway (*11*). This spatial distribution of PI(4,5)P_2_ allows the polar recruitment of proteins to orchestrate key processes, including membrane trafficking (*12–14*) and cytoskeleton remodeling (*15*– *17*). During cytokinesis, while most of anionic lipids accumulate at the cell plate from its inception, PI(4,5)P_2_ is excluded (*18, 19*). Yet, how and when the cell plate changes its identity to become a bona fide, PI(4,5)P_2_-enriched plasma membrane remains unknown.

Here we report that PI(4,5)P_2_ enrichment occurs upon first contact of the growing cell plate with the maternal membrane upon its unilateral attachment. At the same time, PI(4,5)P_2_ becomes excluded from the leading edge of the non-attached side of the phragmoplast, suggesting an active mechanism preventing further passive diffusion. Depletion of PI(4,5)P_2_ at the leading edge correlates with enrichment of the putative PI(4,5)P_2_ phosphatase SAC9. Loss of *sac9* leads to ectopic accumulation of PI(4,5)P_2_ at the leading edge of the phragmoplast, which correlates with MAP65-3 mislocalization and aberrant cell plate branching. We propose a model in which the PI(4,5)P_2_ acts as a spatial cue to guide the leading zone of the phragmoplast at the final step of the plant cytokinesis.

## RESULTS

We investigated how PI(4,5)P_2_ membrane signature is acquired during cytokinesis using the Arabidopsis root meristem as a model. Live cell imaging in four dimensions confirmed that during cytokinesis, PI(4,5)P_2_ biosensors (mCit-TUBBYc or 2mCH-PH^PLC^)(*20*) are excluded from the expanding cell plate until its unilateral attachment to the mother wall (Fig.1a, 1b and Extended data 1, 2. Indeed, at this stage, cell plate attachment is not synchronous, providing an internal control (attached/non-attached on the same cell plate; Fig.1a, blue arrow). These observations suggested that PI(4,5)P_2_-enrichment at the maturing cell plate (Fig. 1b, white arrow) probably arises by diffusion from the highly fluid lateral plasma membrane of the mother cell.

At the leading zone of the same cell plate, where the phragmoplast was not yet attached, significant depletion of the PI(4,5)P_2_ biosensor (index < 1) was observed (Fig.1c to 1f and Extended data 1, 3 suggesting an active mechanism preventing PI(4,5)P_2_ diffusion from the maturing cell plate to the newly formed membrane domain at the cell plate leading zone.

To identify the molecular processes leading to exclusion of PI(4,5)P_2_ from the leading zone in late cytokinesis, we followed the subcellular localization of the plant-specific enzyme SAC9 which participates in the depletion of PI(4,5)P_2_ at the plasma membrane during endocytosis (*11*). We observed that mCit-SAC9 is enriched at the phragmoplast leading zone (index > 1, Fig. 2a to 2c, yellow arrow, Extended data 4, 5, suggesting a role of SAC9 in PI(4,5)P_2_ depletion at the leading zone. To test this hypothesis, we analyzed the localization of a catalytically inactive variant of SAC9 (tdTOM-SAC9^C459A^), for which the signal in the cytoplasm is reduced compared with mCIT-SAC9, allowing imaging at higher resolution (*11*). Like mCit-SAC9, tdTOM-SAC9^C459A^ is enriched at the leading zone while PI(4,5)P_2_ is depleted, leading to a clear mutual exclusion between the enzyme and its substrate along the cell plate (Fig. 2d to 2f, Extended data 4, 6. When SAC9 or the allelic variant SAC9^C459A^ was visualized together with the MAP65-3, a temporal and spatial colocalization at the leading edge of the cell plate was observed, a feature shared by only a few proteins (*4, 8*)(Fig. 3a to 3d, Extended video 1). These results suggest that active restriction of PI(4,5)P_2_ from the leading zone during cell plate attachment is mediated through the enzymatic activity of SAC9.

**Fig.2.**
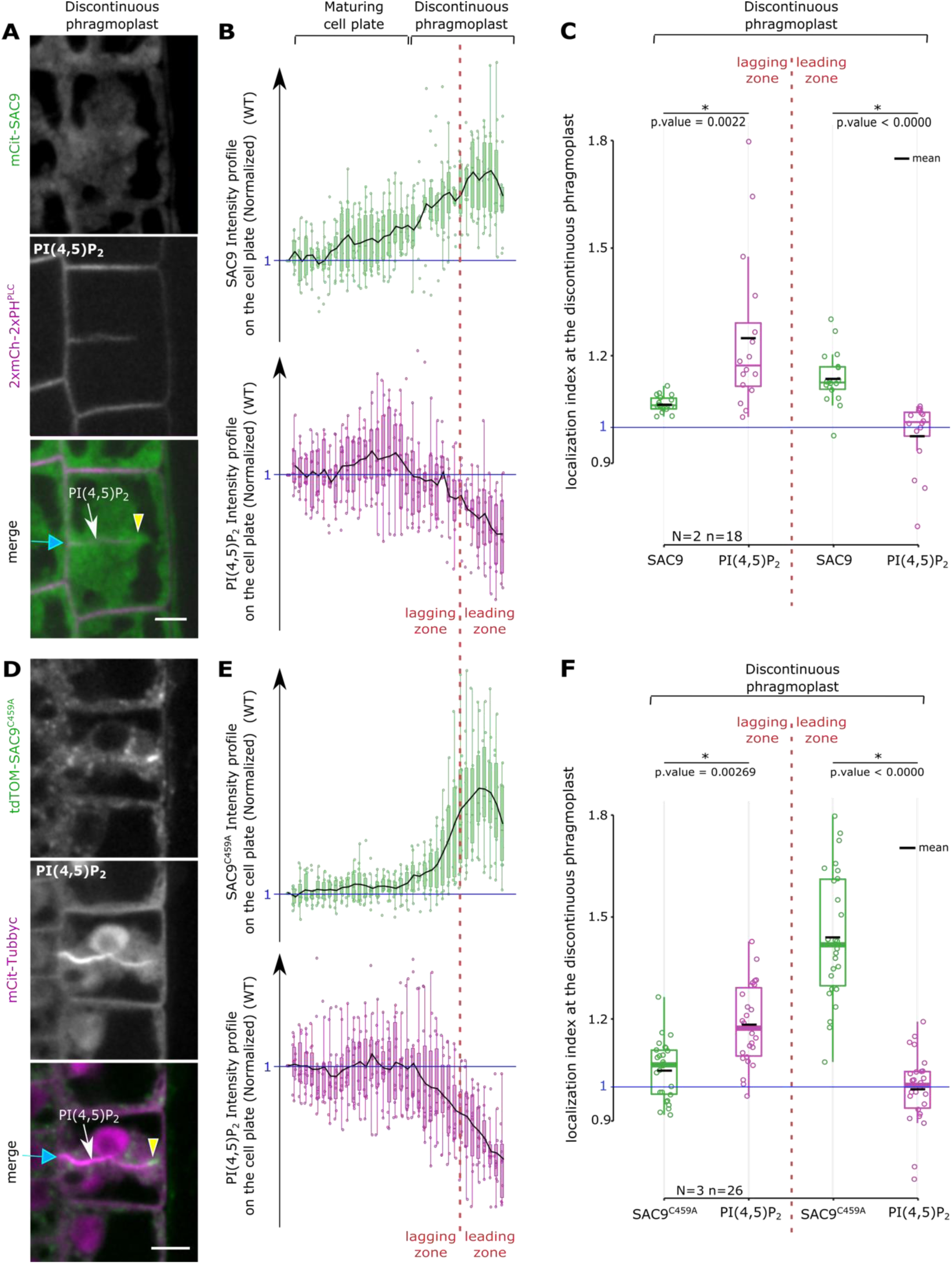
SAC9 and the PI(4,5)P2 are mutually co-excluded at the cell plate. **a**, Images of mCit-SAC9 with 2xmCh-2xPH^PLC^ when the cell plate is unilaterally attached. **b**, normalized intensity profiles corresponding to (a). **c**, Quantification of the localization index corresponding to (a). **d**, Images of tdTOM-SAC9^C459A^ with mCit-Tubbyc when the cell plate is unilaterally attached. **e**, Normalized intensity profiles corresponding to (d). **f**, Quantification of the localization index corresponding to (d). In the plots, middle horizontal bars represent the median, while the bottom and top of each box represent the 25th and 75th percentiles, respectively. At most, the whiskers extend to 1.5 times the interquartile range, excluding data beyond. For range of value under 1,5 IQR, whiskers represent the range of maximum and minimum values. Results of the statistical analysis (shown in the supplementary table) are presented (N = number of replicates, n = number of cells). Blue arrow, cell plate fusion site; yellow arrowhead, SAC9 enrichment; red dotted line, separation leading and lagging zone; n, nucleus; scale bar, 5µm.

If true, the absence of SAC9 should affect PI(4,5)P_2_ distribution at the cell plate. We thus assessed the cytokinetic phenotype of *sac9-3* in which PI(4,5)P_2_ is no longer restricted to the plasma membrane, but also ectopically accumulates in endosomes (*21*). After initial attachment of the cell plate, the PI(4,5)P_2_ biosensor enrichment at the maturing cell plate was observed in *sac9-3* as in the WT (Fig.4a to 4e, white arrow). However, in *sac9-3* we also observed an ectopic accumulation of PI(4,5)P_2_ at the leading edge of the cell plate, probably originating from cytokinetic vesicles (index >1, Fig.4a to 4e, white arrowhead, Extended data 7, 8. Observations in 3D revealed that in *sac9-3*, the PI(4,5)P_2_ biosensor labelled the upfront of the cell plate and then, seems to eventually incorporate to the newly formed plasma membrane during the final attachment (Fig. 4b, Extended data 7, Extended video 2). Moreover, the ectopic accumulation of PI(4,5)P_2_ biosensor at the leading edges of the cell plate observed in *sac9-3* spatially correlates with the accumulation of the non-functional SAC9^C459A^ (Fig. 4f to 4h). The abnormal PI(4,5)P_2_ pattern is consistent with the idea that SAC9 restricts PI(4,5)P_2_ to the cell plate maturing domain and prevents its premature diffusion to the membrane domain formed at the phragmoplast leading zone (Extended data 1, 8).

**Fig.3.**
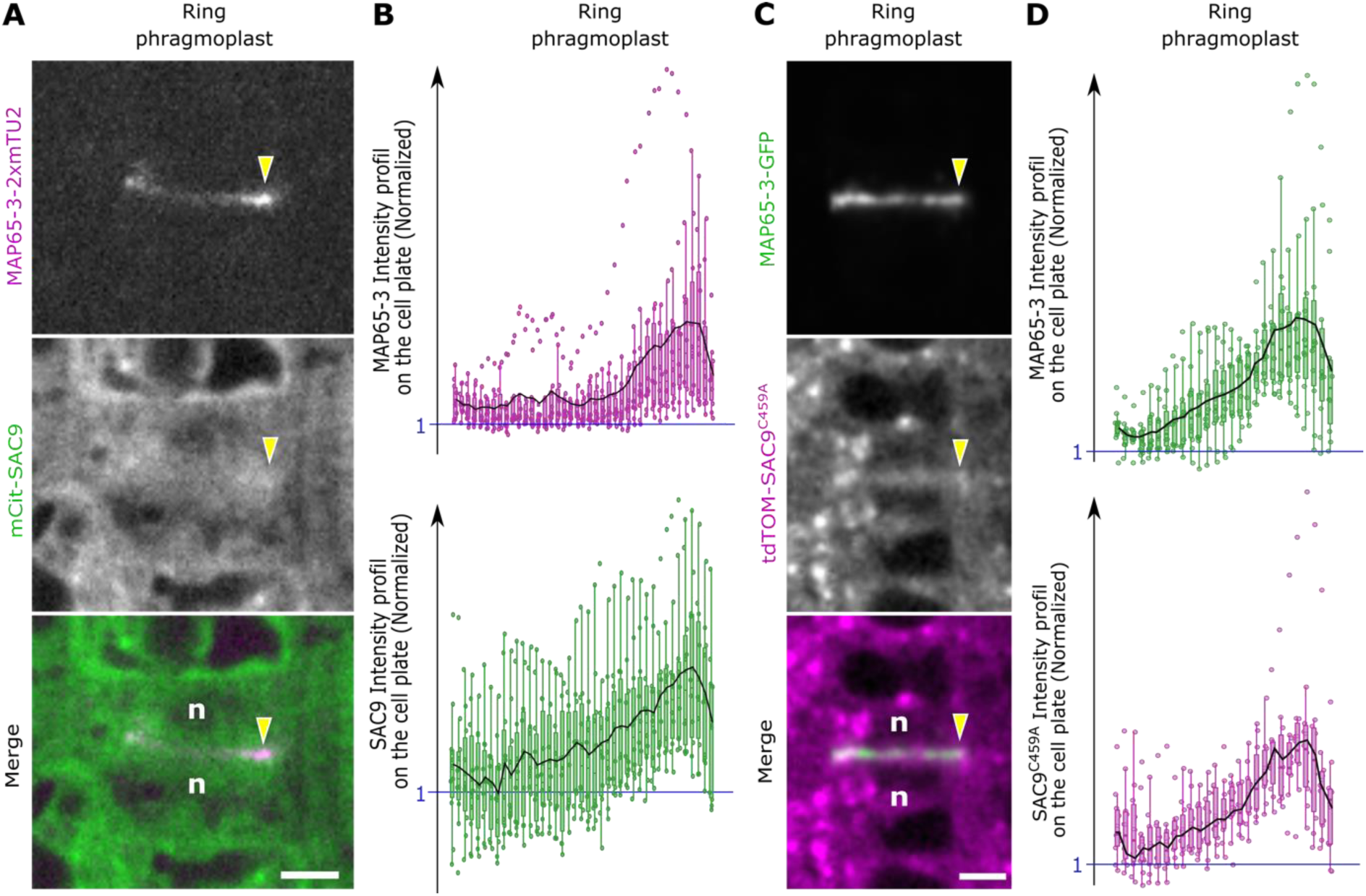
SAC9 colocalizes with MAP65-3 during cytokinesis. **a**, Images of mCit-SAC9 with MAP65-3-2xmTU2. **b**, Normalized intensity profiles corresponding to (a). **c**, Images of tdTOM-SAC9^C459A^ with MAP65-3-GFP. **d**, Normalized intensity profiles corresponding to (c). Red dotted line, separation leading and lagging zone; Yellow arrowhead, SAC9 and MAP65-3 enrichments at the leading zone; n, nucleus; Scale bar, 5 µm.

**Fig.4.**
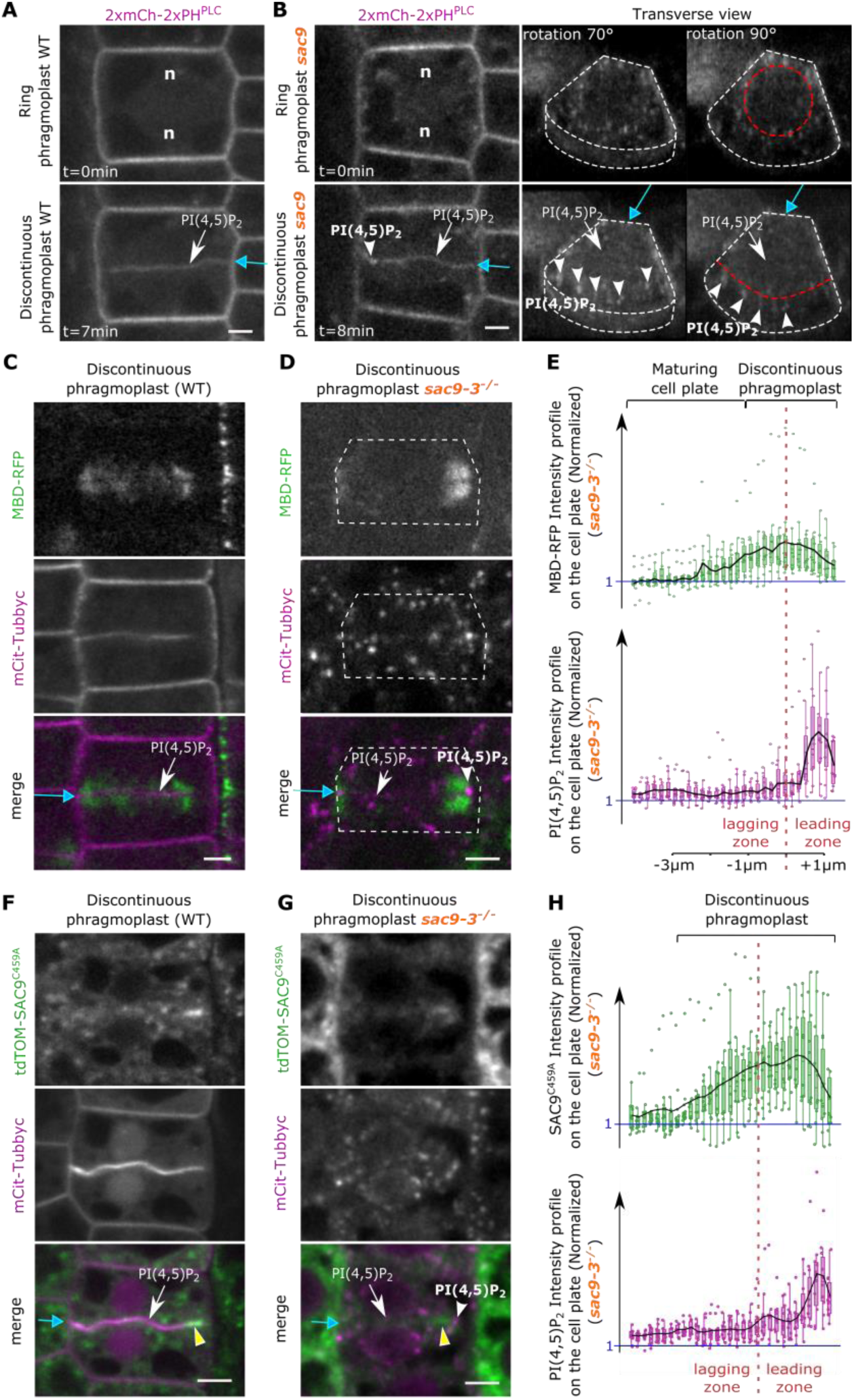
PI(4,5)P2 patterning defects in *sac9-3*. **a**, Images extracted from a timelapse imaging in WT of 2xmCh-2xPH^PLC^ at two cytokinetic steps: ring (upper panel) and discontinuous (lower panel) phragmoplast stages. **b**, Images extracted from a timelapse imaging in *sac9-3* of 2xmCh-2xPH^PLC^ at two cytokinetic steps: ring (upper panel) and discontinuous (lower panel) phragmoplast stages. On the right panel are presented two transverse view of 2xmCh-2xPH^PLC^ for both stages in *sac9-3* (0.3 µm between optical sections). **c**, Images of mCit-Tubbyc with MBD-RFP in WT **d**, Images of mCit-Tubbyc with MBD-RFP in *sac9-3.* **e**, Normalized intensity profiles corresponding to (d). **f** and **g**, Images of the signal corresponding to *tdTOM-SAC9^C459A^* and *mCit-Tubbyc* during the discontinuous phragmoplast phase expressed in a WT (f) or *sac9-3* (g). **h**, normalized intensity profiles corresponding to (g). White arrow, PI(4,5)P_2_ biosensor appearance; White arrowhead, abnormal PI4,5)P_2_ enrichment at the leading zone in *sac9-3*; yellow arrowhead, SAC9^C459A^ enrichment at the leading zone; White dotted line, cell contour; Red dotted line, separation leading and lagging zone; Blue arrow, cell plate fusion site; n, nucleus; Scale bars, 5µm.

Next, we investigated the functional relevance of the SAC9-dependent PI(4,5)P_2_ pattern at the cell plate for cytokinesis. Since phragmoplast organization is regulated by MAP65-3 (*22*–*25*), which colocalizes with SAC9 (Fig. 3a, b), we assessed the localization of MAP65-3 in the absence of SAC9. During ring phragmoplast expansion, MAP65-3-GFP dynamic displays a constant signal width of about 2.06 µm ± 0.37 SD in the WT (Fig. 5a, 5b). The MAP65-3 signal then gradually decreased to eventually disappear upon cell plate attachment. The dynamics of MAP65-3 localization was essentially similar in the *sac9-3* during cell plate expansion. However, in defective cytokinesis observed in *sac9-3*, the MAP65-3-GFP domain was retained at a constant distance of 2.28 µm ± 0.60 SD from the phragmoplast leading edge (Fig. 5a, 5b, Extended video 3, 4, 5), suggesting a failure to transition from the cell plate expansion to its attachment at the cell plate fusion site.

**Fig.5.**
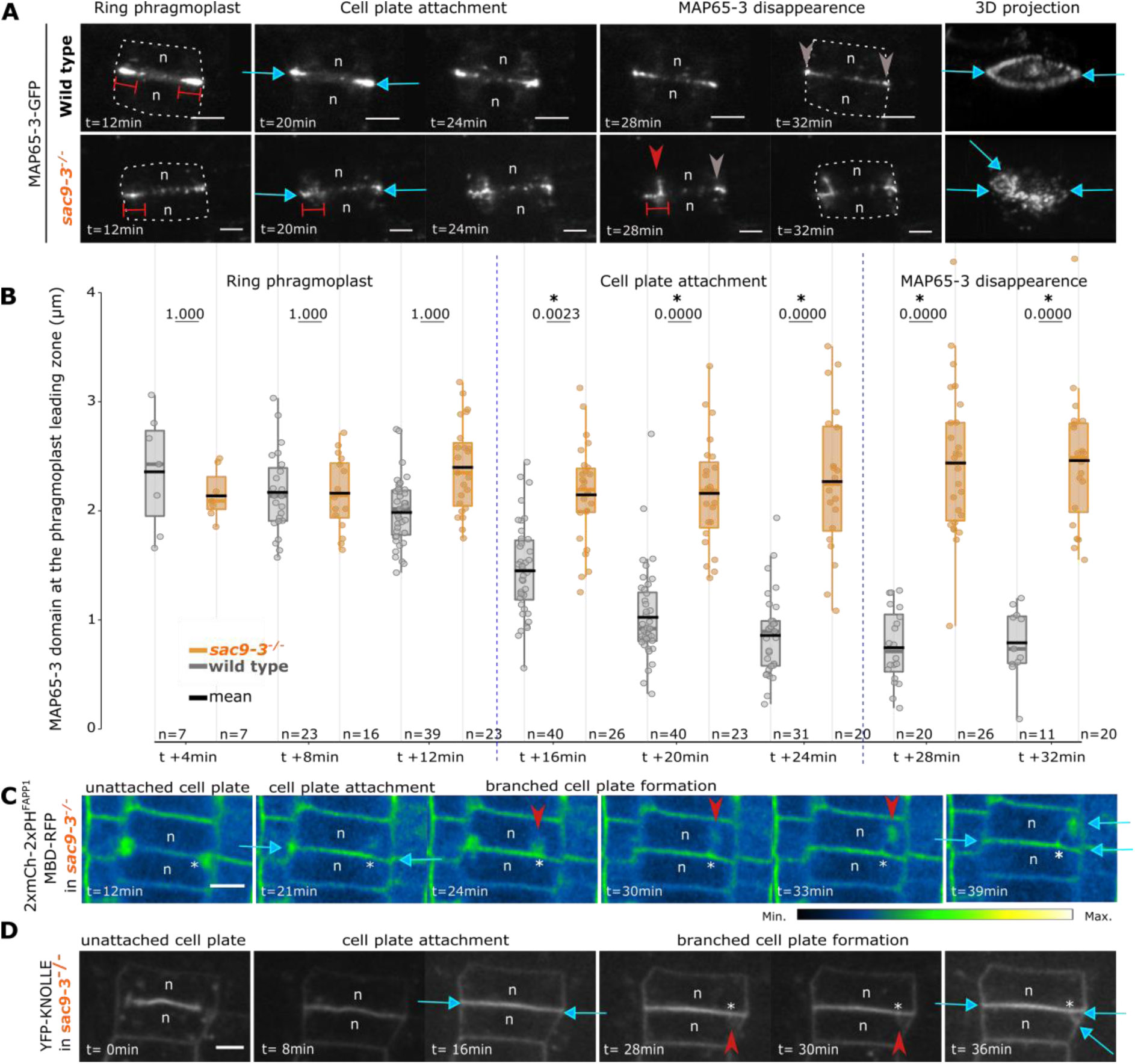
de-novo recruitment of the cytokinesis apparatus in *sac9-3*. **a**, Image series of MAP65-3-GFP in WT (upper panel) or *sac9-3* (of the bottom panel) every two minutes for two hours. Time is given relative to “disk”-“ring” phragmoplast transition (0 min) and cell plate attachment (14 min). Finals images represent a 3D view of time = 32 min. **b**, Comparison of MAP65-3-GFP domain length in WT versus *sac9-3*. In the plots, middle horizontal bars represent the median, while the bottom and top of each box represent the 25^th^ and 75^th^ percentiles, respectively. At most, the whiskers extend to 1.5 times the interquartile range, excluding data beyond. For range of value under 1,5 IQR, whiskers represent the range of maximum and minimum values. Results of the statistical analysis (shown in the supplementary table) are presented (n=number of cells). **c**, Z projection of 2xmCherry-2xPH^FAPP1^ and RFP-MBD in *sac9-3* over time. Here, the signal is color coded in green fire blue (see scale bar on the right). The signal corresponding to microtubules and the one corresponding to membrane are not distinguished). Note that here, mCh-PH^FAPP1^ is used at the membrane marker (plasma membrane and cell plate). **d**, Image series of YFP-KNOLLE in *sac9-3*. Gray arrow, MAP65-3 localization after cytokinesis; Double red bars, distance labeled by GFP-MAP65-3; red arrowhead, branch emergence in *sac9-3*; blue arrow, cell plate fusion site; Asterisk, position of the branch’s emergence at the cell plate; n, nucleus; scale bars, 5µm.

During defective cytokinesis in *sac9-3*, this lack of transition was observed on one side of the cell plate leading zone, and correlated with the emergence of a phragmoplast “branch” from the MAP65-3-GFP inner domain (Fig. 5a, red arrow, Extended data 9a). Eventually, the branch was inserted at an ectopic fusion site (Fig.5a, Extended video 4, 5). On the branch, MAP65-3-GFP localization followed the same pattern as in the main phragmoplast, being progressively restricted to its leading zone and eventually disappearing after the second attachment (Fig.5a, Extended video 4, 5). Observation of the phragmoplast microtubules in *sac9-3* showed that after final cell plate fusion, the phragmoplast disassembled from the main cell plate as it did in the WT (20 min, Fig.5c, Extended data 9b, to 9d). Yet, in aberrant cytokinesis observed in *sac9-3,* a phragmoplast-like structure was reinitiated from the fully expanded cell plate at ∼2-3µm from the fusion site, in a region corresponding to the inner face of the MAP65-3-GFP domain (N= 3, Fig.5a, 5c, Extended data 9b, to 9d).

Since the ultimate effector is the microtubule cytoskeleton, we reasoned that microtubule disruption could lead to a similar phenotype. Treatment of the WT with low concentration of the microtubule inhibitor chlorpropham (*26*) and imaging the membrane marker mCIT-P4M when the different cell files were still organized phenocopied *sac9-3* branched-cell plates, suggesting a link between the phosphoinositides metabolism and the phragmoplast dynamics (N = 3, Extended data 9 extended video 6).

Because in absence of SAC9 the branching emerged from the cell plate (extended video 7) and was decorated by the cell plate marker YFP-KNOLLE (Fig.5d, Fig. 6a,)(*27*), we concluded that this ectopic membrane domain had a cell plate identity. The abnormal branching observed in absence of SAC9 represented 4% of the total apicobasal cells walls (∼ 11 cells / root; n = 30 roots, N = 3 replicates) and was often formed in proximity to the mother cell lateral wall, with an intersection relatively constant at 2.93 µm ± 0.91 SD (Fig. 6b to 6d and Extended data 10, extended video 8, 9), reminiscent of the distance labeled by MAP65-3-GFP and mCIT-SAC9 in WT at the phragmoplast leading zone (Fig. 6b, white asterisk).

**Fig.6.**
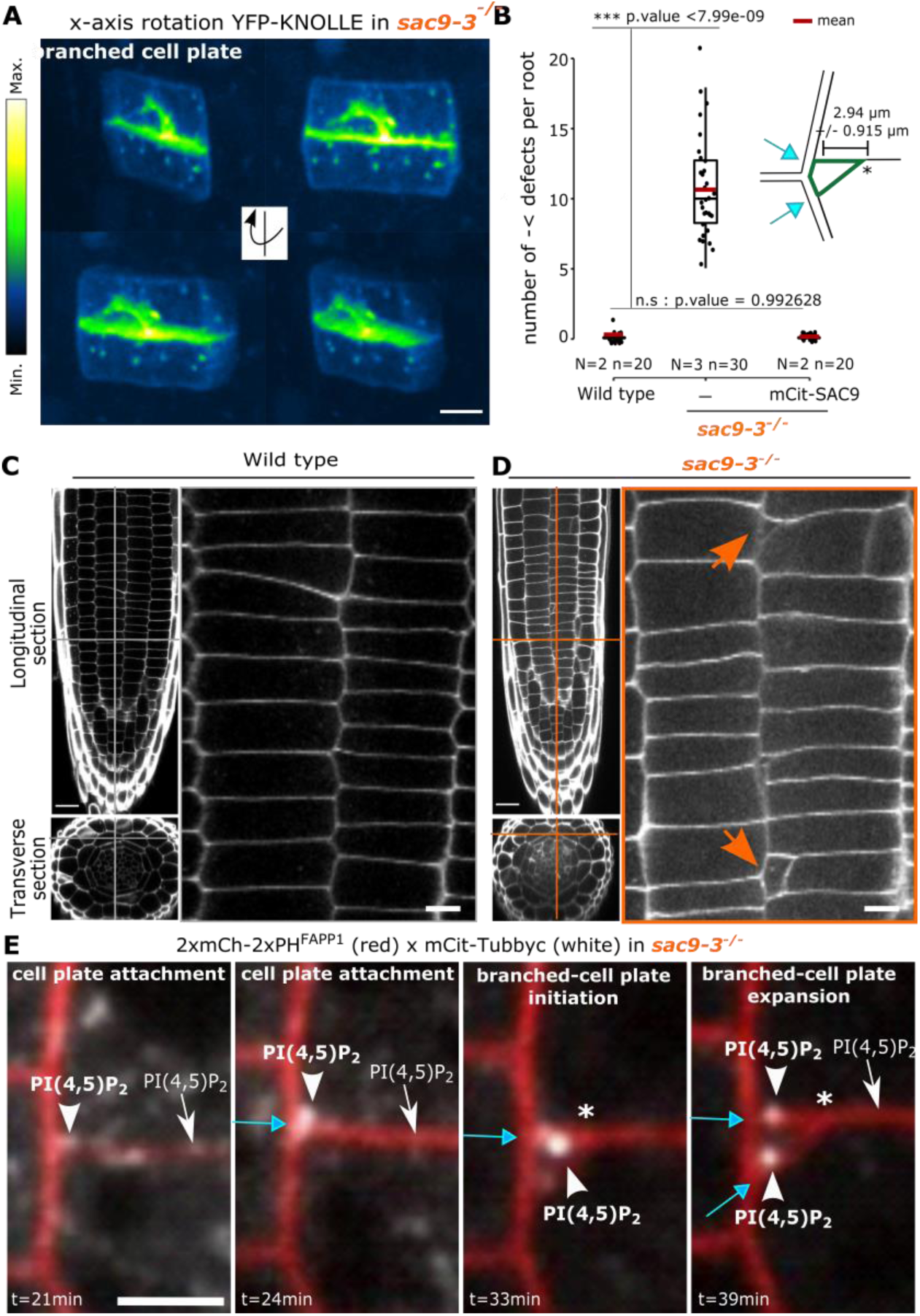
Ectopic membrane domain formed in absence of SAC9 has a cell plate identity and leads to cell wall defects. **a**, 3D rotation of YFP-KNOLLE in *sac9-3* during defective cytokinesis. Here, the signal is color coded in green fire blue (see scale bar). Scale bars, 5µm. **b**, Quantification of the number of defects. The result of the statistical analysis (shown in the supplementary table) is presented. On the top right, representation of branched cell wall defects (green) with key topological elements listed such as the distance between the branched cell wall and lateral cell wall (double arrow), and cell plate insertion site (blue arrow). In the plots, middle horizontal bars represent the median, while the bottom and top of each box represent the 25^th^ and 75^th^ percentiles, respectively. At most, the whiskers extend to 1.5 times the interquartile range, excluding data beyond. For range of value under 1,5 IQR, whiskers represent the range of maximum and minimum values. Results of the statistical analysis (shown in the supplementary table) are presented (N = number of replicates, n = number of cells). **c-d**, Z-stack images of calcofluor-stained fixed roots (7 day-olds) for WT (c) and *sac9-3* (d). Orange lines, the position of the section. Left panel, longitudinal section (top, scale bars 20 µm), and transverse section (bottom). Right panel, a crop of the root cortex (scale bars 5 µm). **e**, Time series of mCit-Tubbyc and 2mCh-2PH^FAPP1^ at the end of the cytokinesis in *sac9-3* (Bars 5µm). Here, 2xmCh-2xPH^FAPP1^ is used as a marker for membranes (plasma membrane and cell plate). n, nucleus; White asterisk, position of the branch’s emergence at the cell plate; Orange arrow, cell wall defects; Blue arrow, cell plate fusion site; White arrowhead, PI(4,5)P_2_ biosensor enrichment observed in *sac9-3*. Scale bars, 5µm.

To test the temporal specificity of this phenotype, we reasoned that a transient rescue may be sufficient. By fusing the regulatory sequence of MAP65-3 (*24*) to SAC9 in order to express SAC9 in mitosis and degrade it after mitosis (Extended data 11), we showed that expressing a functional *SAC9* during mitosis was sufficient to rescue the *sac9-3* cytokinesis defect and part of its dwarf phenotype, highlighting the importance of SAC9 function during cell division.

Concomitantly to abnormal cell plate-branching in *sac9-3*, a specific distribution of PI(4,5)P_2_ was observed, with a burst i) at the cell division site where the cell plate attached, ii) on the cell plate, at ∼ 3 µm where the branching was initiated, iii) and at the ectopic cell division site where the cell plate-branched was inserted (N = 5, white arrowhead, Fig.6e, extended video 10). Moreover, mCIT-*SAC9^C459A^* was not able to rescue the phenotype observed (Extended data 10, 12, suggesting that the phosphatase activity of SAC9 is participating in the observed phenotype.

## DISCUSSION

Cytokinesis in plants determines the topology of cells in a tissue, hence their identities and functions. At the heart of cytokinesis, the coordination between the cytoskeleton and vesicular trafficking is of paramount importance, as it drives the partitioning of the mother cytoplasm into daughter cells. In this context, a variety of proteins implicated in cytoskeleton remodeling and vesicle trafficking have been identified (*2*). Yet, the very final steps of cell division, where the cell plate completes the spatial partitioning of the daughter cytoplasms and acquires the identity of a plasma membrane, remains poorly documented so far.

Here, we assessed the role of membrane anionic lipids during the final steps of cytokinesis. Using Arabidopsis as model, we showed that the anionic lipid signature controls the final step of cytokinesis in plant cells. In the *sac9* mutant, which is impaired in phosphoinositide metabolism(*11*), abnormal PI(4,5)P_2_ signature during the final step of cytokinesis lead to a significant frequency of branched phragmoplasts and multiple cell plate fusion sites in a given cell. Cell plate branching, to our knowledge, is a rare phenomenon that has been only observed after mild perturbation of the microtubule cytoskeleton with drugs or in the conditional mutant *mor1-1* (*28, 29*). This mutant, impaired in a member of the XMAP215 family of microtubule-associated proteins displays at restrictive temperature for 24 h branched cell plates but also incomplete, asymmetric, wandering cell plates (*28*). Even though we cannot conclude that the phenotype observed in *sac9* implies the same mechanism as after microtubule perturbation, the branched-cell plates observed in absence of SAC9 appears therefore more specific than a general disruption of the microtubule cytoskeleton. We wondered if the aberrant branching observed in *sac9* preferentially occurs at the unattached phragmoplast side after unilateral attachment. However, due to the technical limitations of live cell imaging in four dimensions in the native tissue, we cannot answer this point in a quantitative manner. Since the branch is initiated at the existing phragmoplast (and not entirely de novo at random sites), it is reasonable to conclude that affecting microtubule dynamics specifically at the leading edge prior to cell plate fusion induces the initiation of a new dynamic leading edge that originates from the stalled lagging edge.

We found that SAC9 progressively disappears from the fully attached cell plate upon its unilateral attachment to the mother cell. Consequently, SAC9 does not localize to the plasma membrane in the resulting interphasic cells (*11*). SAC9 and its substrate are mutually co-excluded spatially. In sac9, the pool of PI(4,5)P_2_ internalized during defective endocytosis corresponds to a pool present in early endosomes (and probably not to the pool present at the plasma membrane). We, therefore, expect that the internalization of the PI(4,5)P_2_ during defective endocytosis would not be at the expense of the PI(4,5)P_2_ produced at the plasma membrane. It is likely the ratio between the PI(4,5)P_2_ at the plasma membrane and the PI(4,5)P_2_ internalized is abnormal in *sac9-3* rather than the amount of PI(4,5)P_2_ at the plasma membrane. This is likely why the PI(4,5)P_2_ biosensors, in particular, mCIT-Tubbyc, which are expressed at a similar level in WT and *sac9-3*, are mainly labeling the pool of PI(4,5)P_2_ in endosomes than the PI(4,5)P_2_ pool present at the plasma membrane.

In order to test the relationship between the SAC9 localization and function and the patterning of its PI(4,5)P_2_ substrate, we mutated the cysteine in the conserved C-x(5)-R-[T/S] catalytic motif found in all SAC domain-containing phosphoinositide phosphatase (*11*). Here, we showed that while *sac9-3* is complemented with *SAC9pro:mCIT-SAC9*, *SAC9pro:mCIT-SAC9^C459A^*was not sufficient to rescue the cytokinesis phenotypes observed in *sac9-3*. We previously confirmed that mCIT-SAC9^C459A^ fusion was stable and accumulated to a similar extent as mCIT-SAC9 (*11*). Since the probable catalytic cysteine, C459, is required for SAC9 function during cytokinesis, we conclude that the phosphatase activity of SAC9 is necessary to its cytokinetic function. Given that phosphoinositide’s metabolism is highly intricate, we recognize that it is difficult to fully untangle the specific involvement of each lipid in the observed phenotypes. Moreover, SAC9 may carry specific functions outside of its catalytic activity, and therefore the phenotype observed could be due to other factors, such as the recruitment of protein partners.

Given the observed co-localization between SAC9 and MAP65-3, it is tempting to speculate a physical interaction between the two proteins. However, other protein partners, such as RABs, may recruit SAC9 to this location. Indeed, this phenomenon is well known in animal cells, where the functional homologs of SAC9, the PI(4,5)P_2_ 5-phosphatase OCRL is recruited to membranes during cytokinesis via its interaction with RAB35, a member of the RAS Oncogene Family (*30–32*).

We observed that SAC9 undergoes the exact same changes in localization as MAP65-3, first enriched on the entire cell plate (disk phragmoplast stage), then accumulating at the leading edges, and absent from the center of the maturing cell plate. The fact that SAC9 accumulates at the cell plate at an early stage might suggest that PI(4,5)P_2_ accumulation is actively restricted to the expanding phragmoplast. Even though we cannot rule out that SAC9 plays another function during the early phases of phragmoplast formation, the phenotype observed in *sac9* suggests that SAC9 function is required during the cell plate attachment, preventing premature accumulation of the PI(4,5)P_2_ at the growing leading edges.

We show that mitotic expression of SAC9 was sufficient to rescue cytokinetic defects in the *sac9* mutant. This result suggests that restoring the PI(4,5)P_2_ pattern in dividing cells is sufficient to drive proper transition from a maturing cell plate to a plasma membrane identity. MAP65-3^Cter^-SAC9 is also observed in the cytoplasm of the dividing cell, we therefore cannot conclude if the effects is direct. Moreover, since the dwarfism observed in absence of SAC9 is (i) only partially rescued by the expression of *MAP65-3^Cter^-SAC9*, particularly in the aerial part, we cannot rule out that the PI(4,5)P_2_ signature is not fully restored in interphasic cells.

Using PI(4,5)P_2_ biosensors, we observed that PI(4,5)P_2_ is depleted from the leading edge of the cell plate, where SAC9 activity restricts its concentration. Local changes in PI(4,5)P_2_ abundance may provoke modifications in microtubule dynamics and stop phragmoplast expansion. It was recently shown that upon PI(4,5)P_2_ depletion using an inducible system, highly anisotropic transverse cortical microtubules were replaced by randomly arranged filaments, suggesting a function of PI(4,5)P_2_ in the organization of microtubules in plant cells (*15*). Active exclusion of PI(4,5)P_2_ at the unattached leading end of the phragmoplast may thus be required to maintain microtubule dynamics until the successful completion of cytokinesis.

We propose that upon unilateral cell plate attachment to the plasma membrane, polarity domains (i; e. cell plate maturing domain versus cell plate leading edges) are formed and the phragmoplast lagging zone acts as a barrier for PI(4,5)P_2_ diffusion on the cell plate leading edge. Concomitantly to the final cell plate attachment to the mother cell, PI(4,5)P_2_ gradually invades the entire plasma membrane precursor from the maturing zone to the decreasing leading zone. In absence of SAC9, PI(4,5)P_2_ gradient at the unilaterally attached cell plate is in the opposite direction, as PI(4,5)P_2_ biosensor abnormally accumulates at the leading zone. This inverted PI(4,5)P_2_ gradient might perturb the function of proteins such as MAP65-3, directing to the reassembly of the phragmoplast apparatus on the inner side of the leading zone. There, PI(4,5)P_2_ might act as a spatial cue to guide the leading zone of the phragmoplast at the final step of the plant cytokinesis (Extended data 13).

## ACKNOWLEDGEMENTS

We are grateful to the SiCE group in particular Yvon Jaillais and Vincent Bayle for comments and discussions, as well as Charlotte Kirchhelle, Liam Elliot, Olivier Hamant (RDP, ENS de Lyon, France) for their great input on the manuscript. We thank Patrice Bolland, Alexis Lacroix from our plant facility, and Claire Lionnet for microscopy advice. We acknowledge the contribution of SFR Biosciences (UMS3444/CNRS, US8/Inserm, ENS de Lyon, UCBL) facilities at the LBI-PLATIM-MICROSCOPY for assistance with imaging. This work was supported by Seed Fund ENS LYON; ANR-16-CE13-0021 to MCC and ANR-20-CE13-0026-02 to MCC and DB. AL is funded by Ph.D. fellowships from the French Ministry of Research and Higher Education.

## Author contributions

Conceptualization: AL, MCC

Methodology: AL, KB, AF, EG, CG

Investigation: AL, EG, MCC, CG

Funding acquisition: MCC, DB, AL

Supervision: MCC, MP, DB

Writing – original draft: AL, MCC

Writing – review & editing: AL, MCC, MP

## Data and materials availability

All data and materials are available upon request

## Supplementary Materials

Materials and Methods

Figs. S1 to S13

Tables S1 to S10

References (*47*)

Movies S1 to S10

## Materials and Methods

### Growth condition and plant materials

#### Arabidopsis thaliana

Columbia-0 (Col-0) accession was used as a wild-type (WT) reference genomic background throughout this study. Arabidopsis seedlings were grown in vitro on half Murashige and Skoog (½ MS) basal medium supplemented with 0.7% plant agar (pH 5.7) in continuous light conditions at 21 °C. seedlings were imaged between 5 and 7dag and finally grown in soil under long-day conditions at 21 °C and 70% humidity 16 h daylight. For Isopropyl N-(3-chlorophenyl)-carbamate (chlorpropham, PESTANAL^®^, analytical standard) treatments, seedlings grown in vitro for 5 days were then transferred on ½ MS basal medium supplemented with 0.7% plant agar and 20µM of chlorpropham. Images were taken after 2h of incubation.

### Cloning and plant transformation

#### Cloning of MAP65-3pro

*2xmTU2-MAP65-3,* The *MAP65-3pro* (1,2 kb) was flanked with attB2R and attB3 sequences and recombined by BP gateway reaction into pDONR221 (Fw: GGGGACAACTTTGTATAGAAAAGTTGCTTACACTCTTCCCTACACAAAACCGCG ; Rv: GGGGACTGCTTTTTTGTACAAACTTGCTTCGAAATGCTTAAGCCTGTAACAGG). Final destination vectors (MAP65-3pro/P5’, MAP65-3/pDON207 (*33*), 2xmTU2/pDONR-P3’) were obtained using three fragments LR recombination system (life technologies, www.lifetechnologies.com/) using pK7m34GW destination vector (*34*).

The collection of phosphoinositide biosensors used in this study was described previously(*20*). A tandem dimer of the monomeric red fluorescent protein CHERRY (2xmCH)(*35*) was used to generate a stable transgenic Arabidopsis line to increase the poor signal obtain in red, in particular, because of the autofluorescence of the plant tissue.

The swapping domain was engineered using a vector containing mCitrine, cYFPnoSTOP/pDONR221 (previously described by (*20*)), and a vector containing MAP65-3 (*33*) . The C-terminal domain 2 of MAP65-3 (MAP65-3^Cter^; EALYGSKPSPSKPLGGKKAPRMSTGGASNRRLSLGAAMHQTPKPNKKADHRHNDGALSNGRRGLDIAGLPSRKQSMNPSEMLQSPLVRKPFSPISTTVVASKANIATTTTQQLPKNNAVNEISSFATPIKNNNILRNLEEEKMMTMMMQTPKNVAAMIPIPSTPATVSVPMHTAPTPFTNNARLMSEKPEVVEYSFEERRLAFMLQSECRLV) was amplified by PCR (*36*): FW: ccaactttgtacaaaaaagcaggctttaaccatggaggcactttacgggtccaaaccc; Rv: gaacagctcctcgcccttgctcaccataaccaaacgacattcagactgtagcatgaa) and introgressed into cYFPnoSTOP/pDONR221 by Gibson cloning (primer FW: ccttcatgctacagtctgaatgtcgtttggtaatggtgagcaagggcgaggagctgt Rv: gctgggtttggacccgtaaagtgcctccatggttaagcctgcttttttgtacaaagtt). Then by directed mutagenesis we add an ATG before MAP65-3^Cter^ sequence (primer FW:gcaggcttaaccATGgaggcactttacggg Rv:cccgtaaagtgcctcCATggttaagcctgc). MAP65-3^Cter^ -cYFPnoSTOP/pDONR221vector was used to obtain final destination vectors using three fragments LR recombination system (life technologies, www.lifetechnologies.com/) using pB7m34GW destination vector (*34*), pDONR-P4-P1R containing MAP65-3 promotor and pDONOR-P2R-P3 containing SAC9.

Wild-type Col-0 and heterozygous (or homozygous) *sac9-3*were transformed using the dipping method (*37*). For each construct generated in this paper, between 20 and 24 independent T1 were selected on antibiotics and propagated. In T2, all lines were screened using confocal microscopy for fluorescence signal and localization. From three to five independent lines with a mono-insertion and showing a consistent, representative expression level and localization were selected and grown to the next generation. Each selected line was reanalyzed in T3 by confocal microscopy to confirm the results obtained in T2 and to select homozygous plants. At this stage, we selected one representative line for in depth analysis of the localization and crosses and two representative lines for in depth analysis of mutant complementation.

### Live cell imaging

Time-lapse imaging on living root tissues were acquired either manually, or using an automated root tracking system set-up previously (*38*). Briefly, plants roots expressing fluorescent proteins were imaged with a spinning disk confocal microscope while they grow using automatic movement of the microscope stage that compensates for root growth, allowing the imaging of the dividing cells in the root meristem over time (*38*). For both type of experiment, Z-stacks were acquired every 1 min to 3 min with the following spinning disk confocal microscope set up: inverted Zeiss microscope (AxioObserver Z1, Carl Zeiss Group, http://www.zeiss.com/) equipped with a spinning disk module (CSU-W1-T3, Yokogawa, www.yokogawa.com) and a ProEM+ 1024B camera (Princeton Instrument, www.princetoninstruments.com/) or Camera Prime 95B (www.photometrics.com) using a 63x Plan-Apochromat objective (numerical aperture 1.4, oil immersion). GFP and mCITRINE were excited with a 488 nm laser (150 mW) and fluorescence emission was filtered by a 525/50 nm BrightLine! a single-band bandpass filter (Semrock, http://www.semrock.com/), mCHERRY and TdTOM were excited with a 561 nm laser (80 mW) and fluorescence emission was filtered by a 609/54 nm BrightLine! a single-band bandpass filter (Semrock). For quantitative imaging, pictures of epidermal root meristem cells were taken with detector settings optimized for low background and no pixel saturation. Care was taken to use similar confocal settings when comparing fluorescence intensity or for quantification. In this study, we used the PI4P biosensor (FAPP1) as a general marker for endomembrane as it localizes both to plasma membrane and Trans-Golgi Network (*18*). This allows the visualization of both the plasma membrane and the cell plate (*18*), in *sac9-3* and WT plants. We are not expecting a change in behavior for this biosensor in sac9 as we observed the same localization in the WT (*11*). For the double localization of the membranes and the microtubules in *sac9-3* using time-lapse imaging (Figure 5c and Extended data 9d), the fact that both markers are tagged with similar fluorophores is not optimal. In fact, we were only able to rescue plants for the cross *sac9-3^-/-^* x MBD-RFP. Then we tried to introduce a third construct in *sac9-3^-/-^* x MBD-RFP with a mCIT tagged membrane marker, but we did not succeed. Therefore, because of technical limitations, we analyzed the line expressing sac9-3^-/-^ x MBD-RFP x 2xmCherry-2xPH^FAPP1^ and the corresponding MBD-RFP x 2xmCherry-2xPH^FAPP1^ in WT.

### Calcofluor staining and immunolocalization imaging

For calcofluor staining, root meristem cell walls were stained using the calcofluor dye, following the protocol described in (*39*). Seedlings were incubated overnight in a fixation buffer (50% of methanol, 10% of acetic acid, and 40% of distilled water). The seedlings were then rehydrated in ethanol baths for 10 min each: 50% ethanol, 30% ethanol, and distilled water twice. Afterward, the seedlings were transferred in the staining solution for overnight incubation (90% of clearsee solution (5% of urea, 15% of deoxycholic acid, and 10% of xylitol in distilled water) and 10% of calcofluor white solution (500 mg Fluorescent Brightener 28 in distilled water (qsp 50mL) and 1 drop of NaOH 10N). Before imaging, the seedlings were rinsed for 15 min in clearsee solution. For segmentation and tissue localization of the defects, z-stacks were performed with 0.39 µm space between acquisitions. For *sac9-3* defects comparison with MAP65-3^Cter^ complementation experiments, z-stacks were performed with 0.8 to 1µm space between acquisitions.

Whole-mount immunolocalization was performed as described (*40*). For immunolocalization, seedlings were fixed in 4% paraformaldehyde and 0.1% Triton X-100 in ½ MTSB buffer (25 mM PIPES, 2.5 mM MgSO4, 2.5 mM EGTA, pH 6.9) for 1 h under vacuum, then rinsed in PBS 1X for 10 min. Samples were then permeabilized in ethanol for 10 min and rehydrated in PBS for 10 min. Cell walls were digested using the following buffer for 1 h: 2 mM MES pH 5, 0.20% driselase and 0.15% macerozyme. Tissues were incubated overnight at room temperature with the B-5-1-2 monoclonal anti-α-tubulin (Sigma) and the anti-KNOLLE antibody (kind gift of G. Jürgens, University of Tübingen, Germany (*41*)). The next day, tissues were washed for 15 min in PBS, 50 mM glycine, incubated with secondary antibodies (Alexa Fluor 555 goat anti-rabbit for KNOLLE antibody and Alexa Fluor 488 goat anti-mouse for the tubulin antibody) overnight and washed again in PBS, 50 mM glycine. Samples were incubated in 10% calcofluor white solution for 2 hours and then mounted in VECTASHIELD.

For imaging mCIT-SAC9 in fixed tissue, seedlings expressing mCIT-SAC9 were fixed in 4% paraformaldehyde and 0.1% Triton X-100 in ½ MTSB buffer (25 mM PIPES, 2.5 mM MgSO4, 2.5 mM EGTA, pH 6.9) for 1 h under vacuum, then rinsed in PBS 1X for 10 min. Samples were then permeabilized in ethanol for 10 min and rehydrated in PBS for 10 min. Cell walls were digested using the following buffer for 1 h: 2 mM MES pH 5, 0.20% driselase and 0.15% macerozyme. Tissues with preserved fluorescence were mounted in VECTASHIELD and viewed using an inverted Zeiss CLSM800 confocal microscope using a 40x Plan-apochromatic objective. Dual-color images were acquired by sequential line switching, allowing the separation of channels by both excitation and emission. GFP was excited with a 488 nm laser, mCIT was excited with a 515nm laser, mCH/tdTOM were excited with a 561 nm laser and finally, Fluorescent Brightener 28 (Calcofluor) was recorded using a 405 nm excitation.

### Segmentation and tissue localization of *sac9-3* defects

Quantification of cell division defects was based on manual counting of full Z-stacks with a z-spacing equal to the lateral resolution in order to get cubic voxels allowing to quantify the defects, number, and position in the tissue, with the ImageJ orthogonal view plugin (or z-spacing of 0.5 and 1µm in cases that only defects count was needed). Segmentation was performed using ImageJ, (morphological segmentation) plugin as described and angle extraction using ImageJ plugin developed (*42*).

The *sac9-3* defects architecture was manually measured using ImageJ, “straight line” and “angle tool” tools, and data were analyzed with excel.

### Fluorescence intensity extraction and localization indexes

<u>Normalized fluorescence intensity</u>: was obtained using Fiji tool, extracted from 10 dividing cells (cells of an approximatively equivalent length, normalized to the exact same size by suppressing a part of the space between the cell plate and plasma membrane (Disk phragmoplast) or just the exact same length on only one expanding edge (discontinuous phragmoplast), normalized by the intensity at the cell plate center (except for Fig3, Extended data 4 and 6 where the normalization was by the cytosolic intensity), and plotted to obtain an average intensity profile.

<u>mCit-SAC9 fluorescence intensity extraction:</u> For images obtained in fixed samples, the fluorescence intensity was obtained using Fiji tool. Three-axis: cell plate axis, across the center of the cell (in the root growth axis), and finally across the cell cortex (nearby the lateral cell wall in the root growth axis) were drawn through the cell and the signal intensity was plotted using the Plot Profile tool.

<u>For PI(4,5)P_2_ localization index on the discontinuous phragmoplast:</u> phragmoplast length was measured and divided into two equal parts, the leading zone (domain close to the plasma membrane) and the lagging zone (domain close to the cell center). Fluorescence intensity was measured on each zone of the phragmoplast and, in addition, on the entire cell plate and cytosol. Next, the cytosol intensity background was subtracted and, finally, the leading and lagging zones were divided by the global cell plate intensity. The final ratio superior to 1 underlined an enrichment at a specific zone of the phragmoplast. The final ratio inferior to 1 underlines the depletion.

<u>SAC9-PI(4,5)P_2_ localization index on the discontinuous phragmoplast:</u> To address mCit-SAC9 and tdTOM-SAC9^C459A^ co-exclusion with PI(4,5)P2 biosensor we used the same index as the PI(4,5)P_2_ localization index. Because we showed that the PI(4,5)P_2_ biosensor was excluded from the leading zone, when a microtubule marker was not present in the imaged transgenic line, we decided to use the PI(4,5)P_2_ biosensor as a separation between the leading and lagging zone. Using PI(4,5)P_2_ biosensor fluorescence decrease as a marker of this separation, we measured the intensity on 1.5 µm (the approximative half-length of the phragmoplast) on each side of PI(4,5)P_2_ biosensor limit. Finally, the leading and lagging zones were divided by the intensity of the cell plate and nearby cytoplasm (1µm around the cell plate).

<u>MAP65-3 labeling distance:</u> Measurement was performed using ImageJ, a “straight line” tool every two minutes on one side of the cell plate (the one with the persistent pool in *sac9-3*). Lines were created between the outer and inner MAP65-3 on the entire length labeled. The lengths were plotted relative to the “disk”-“ring” phragmoplast transition (0 min) and cell plate attachment (14 min).

<u>PI(4,5)P_2_ phragmoplast accumulation index</u>: One dividing cell and one of its surrounding non-dividing cells were taken for each root. For every two cells, the signal intensity was measured in one elliptical region of interest (ROIs) at the phragmoplast (around the cell plate across all cell lengths or at the equator of interphasic cells), with an equal area between the two cells. In addition, we measured signal intensity at the plasma membrane separating the two cells. Ratios were obtained by dividing both elliptical regions by the signal intensity at the plasma membrane (obtaining the “dividing ratio” for the dividing cell and the “non-dividing ratio” for the non-dividing cell). Finally, the ratio between the “dividing ratio” and the “non-dividing ratio” was analyzed. Final ratios equal to 1 indicated that there was no enrichment during cell division. Final ratio greater than 1 indicated an enrichment during cell division at the phragmoplast. Final ratio less than 1 indicated a depletion during cell division at the phragmoplast. This index was performed on the young dividing cells where the cell plate is expanding and not attached and in old dividing cells where the cell plate is partially attached to the plasma membrane (discontinuous phragmoplast).

<u>MAP65-3 accumulation at one side of the cell plate index:</u> This index was used to interpret MAP65-3-GFP accumulation on the edges of the cell plate. Using ImageJ, we measured fluorescence intensity on the cell plate edges observed in 2 dimensions at Ziii (at the Z position where the cell plate is the most expanded, Extended data 9a) and at the center of the cell plate (equivalent ROIs designed to feet to the left side in WT or the side where the defect is formed in *sac9* mutant). Then fluorescence intensity values at the cell plate leading zone were divided by the intensity value at the center of the cell plate and finally, we calculated the ratio between the intensity value at the defect side/center (for wild type plants we randomly defined the left edge of the cell plate to make the analysis) and the intensity value at the second (right cell plate edge for wild type plants) side/center. Index strongly superior to 1 highlights a polarized accumulation of MAP65-3 on one zone of the cell plate edges in late cell division.

### Statistical analysis

We performed all our statistical analyses in R (version 3.5.0 (2018-04-23), using R studio interface and the packages ggplot2 (*43*), lme4 (*44*), car (*45*), multcomp (*46*) and means (*47*). Graphs were obtained with R and R-studio software, and customized with Inkscape (https://inkscape.org).

The number of division defects was compared using a generalized linear model with a Poisson law.

Fluorescence intensity indexes were compared using an ANOVA statistical test and a Thuckey HSD post hoc analysis if the normality was verified or using a Kruskal-Wallis test, Dunn (Bonferroni method) Post hoc analyses, if not.

## SUPPLEMENTARY FIGURES

**Extended data.1.**
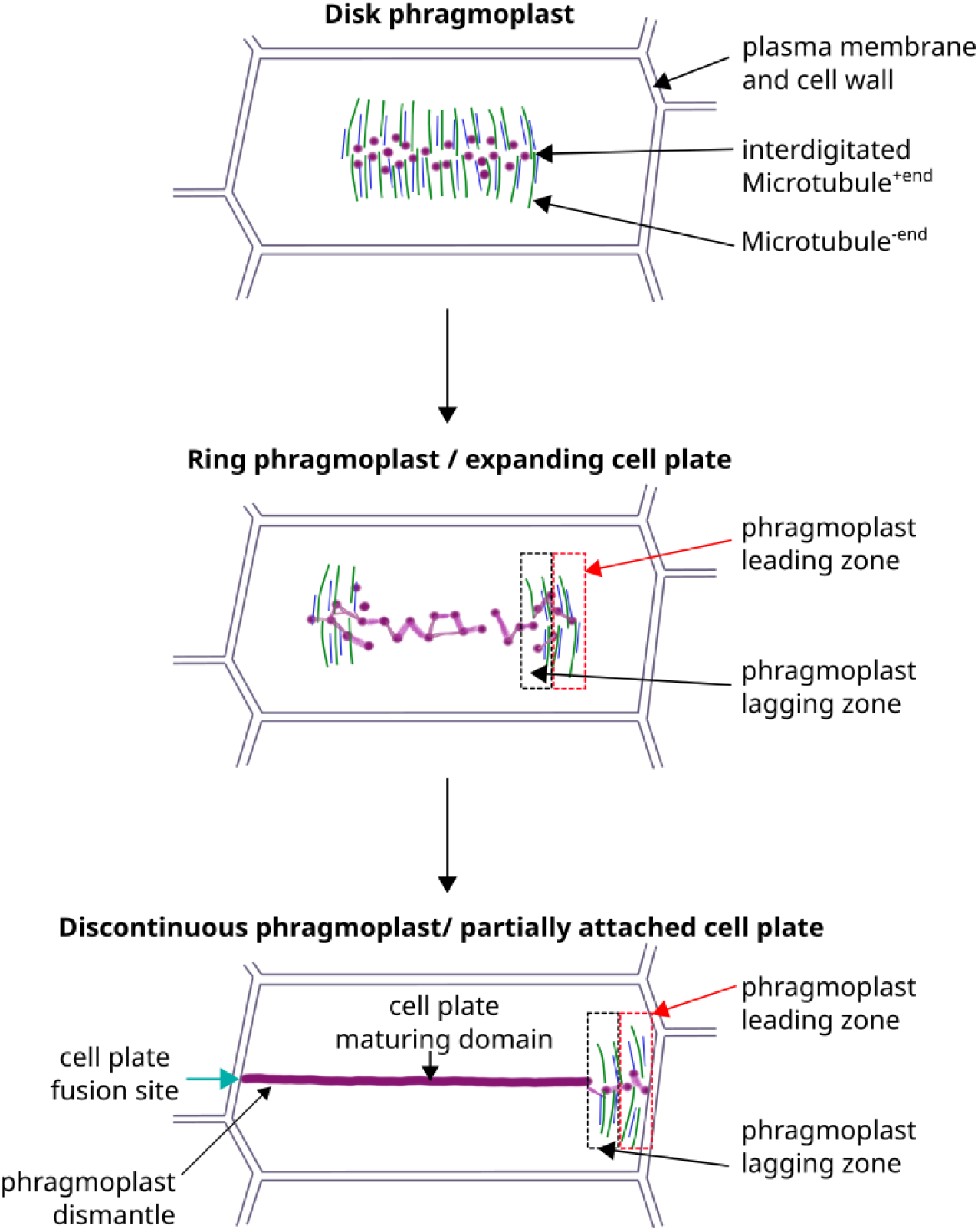
Cell cytokinesis from a cytoskeleton point of view. Schematic representation of phragmoplast expansion in a 2D cell. The three main phragmoplast structures are presented: form a Disk phragmoplast until a Discontinuous phragmoplast morphology. Key cytoskeleton’s structure are presented: Microtubules plus ends, minus ends, phragmoplast leading and lagging zones and cell plate maturing domain. Microtubules, green; Actin, dark blue, Cell plate, magenta.

**Extended data.2.**
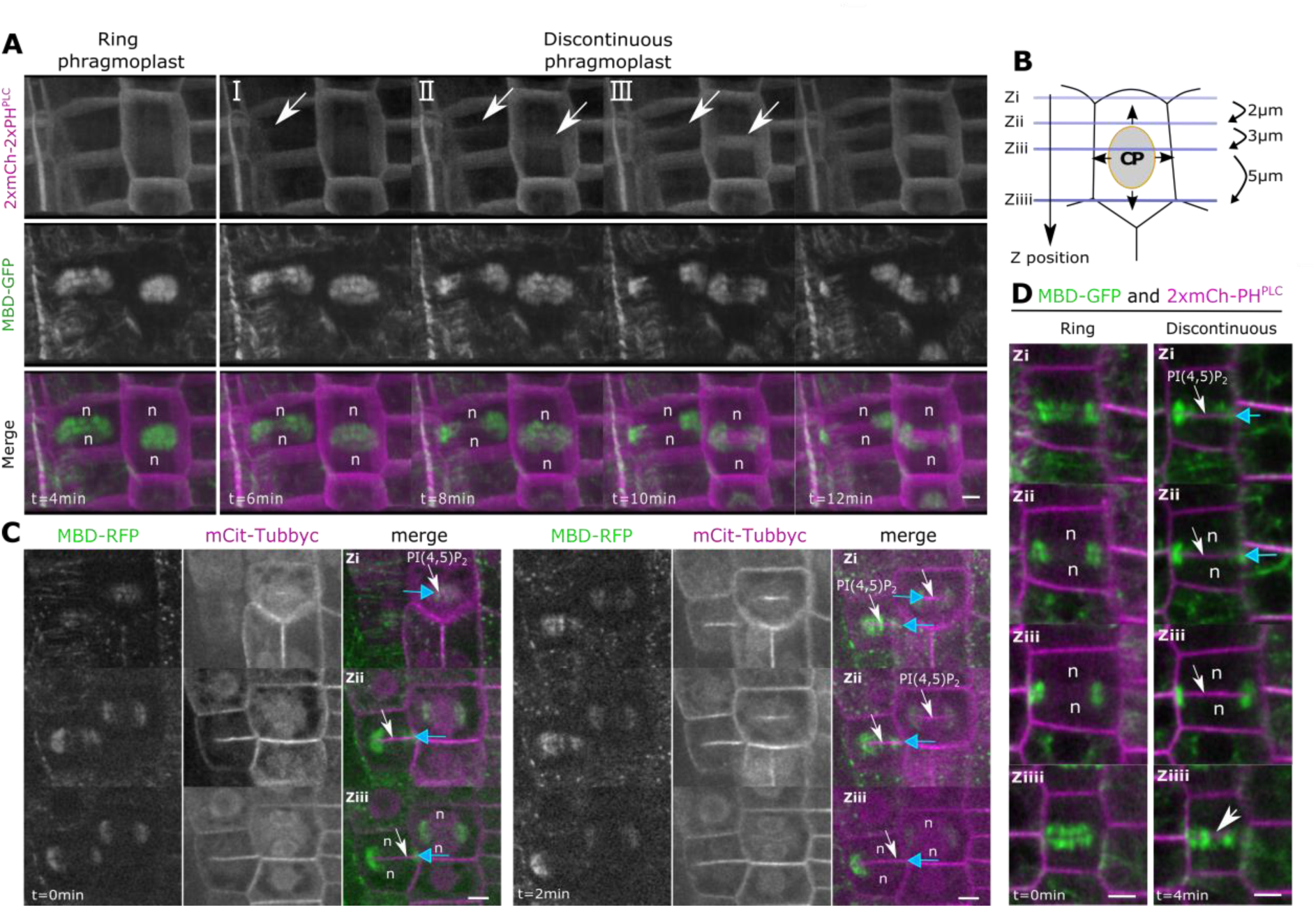
PI(4,5)P_2_ appearance at the maturing cell plate in wild-type plants. **a**, Z-projection of dividing cells expressing 2xmCh-2xPH^PLC^ together with MBD-GFP. The image was taken every 2 min and 15 optical sections were used for Z-projection. I, II and III images correspond to the sequence of events from the first appearance of PI(4,5)P_2_ biosensor at the cell plate. **b**, Representation of the different focal planes used in this study from the subcortical part of the cells (Zi, top panel) to the bottom cells (Ziiii, bottom panel). **c**, Confocal images of WT at different Z, expressing *mCit-Tubbyc* together with *MBD-RFP* during the discontinuous phragmoplast stage, in wild type plants. **d**, Confocal images of WT, expressing *2xmCh-2xPH^PLC^* together with *MBD-GFP* during the discontinuous phragmoplast stage, from the subcortical part of the cells (Zi, top panel) to the center of the cells (Ziiii, bottom panel) in WT. White arrow, PI(4,5)P_2_ biosensor appearance; blue arrow, cell plate fusion site; Black arrow, cell plate direction of growth; n, nucleus; scale bars, 5µm.

**Extended data.3.**
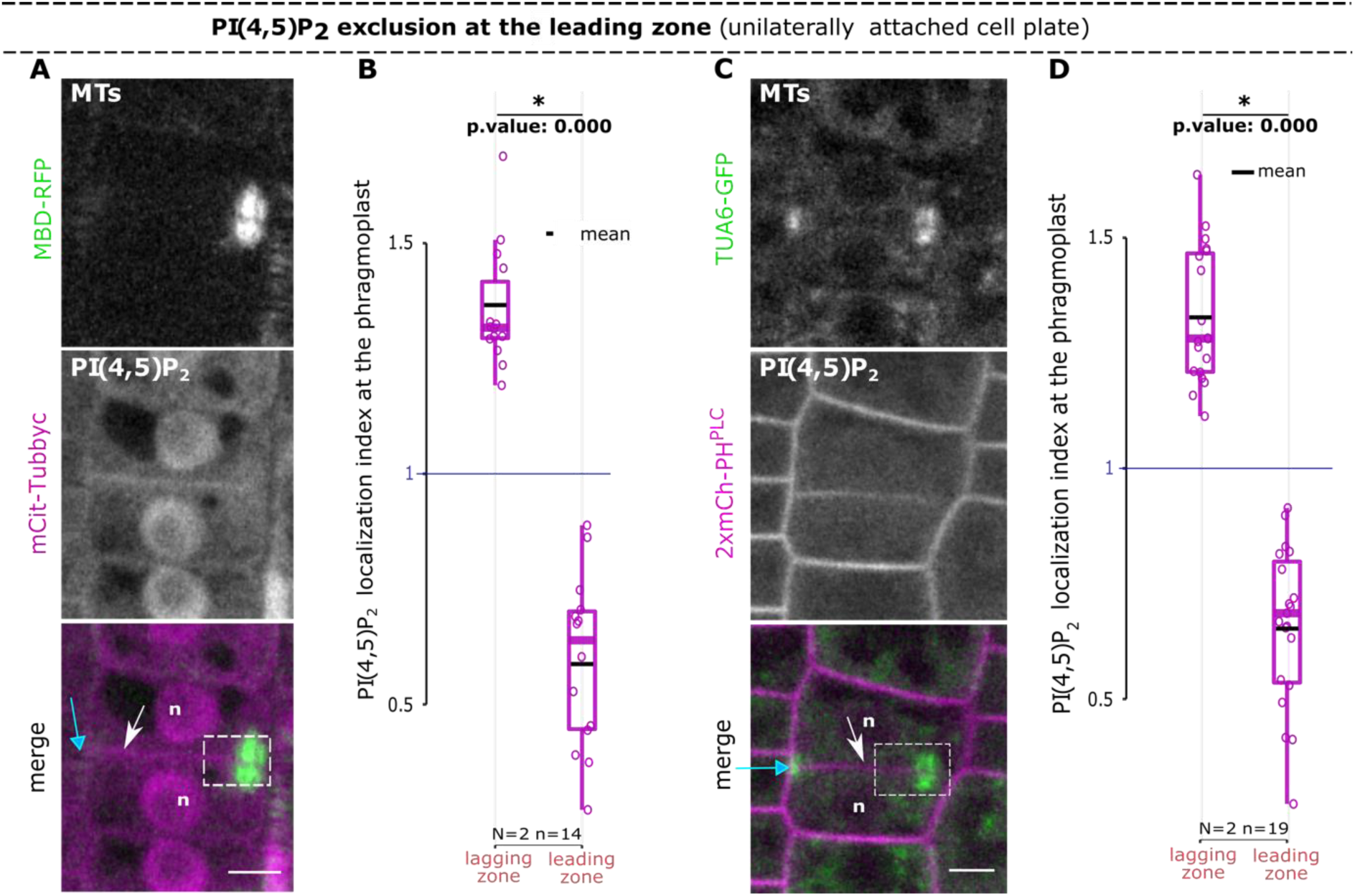
PI(4,5)P_2_ exclusion at the leading zone. **a, c**, Confocal images of WT, expressing respectively *mCit-Tubbyc* with *MBD-RFP* (a) or *2xmCh-2xPH^PLC^* with *TUA6-GFP* (c) at the discontinuous phragmoplast. **b, d**, Quantification of PI(4,5)P_2_ localization index for PI(4,5)P_2_ biosensors (mCit-Tubbyc and 2xmCh-2xPH^PLC^) with respect to two microtubules tagged proteins (MBD-RFP and TUA6-GFP respectively), normalized by cell plate intensity. “1” blue line represents a situation in which there is no biosensor enrichment. In the plots, middle horizontal bars represent the median, while the bottom and top of each box represent the 25^th^ and 75^th^ percentiles, respectively. At most, the whiskers extend to 1.5 times the interquartile range, excluding data beyond. For range of value under 1,5 IQR, whiskers represent the range of maximum and minimum values. Results of the statistical analysis (shown in the supplementary table) are presented (N = number of replicates, n = number of cells). White arrow, PI(4,5)P_2_ biosensor appearance; Blue arrow, cell plate fusion site; dotted lines, region of interest; n, nucleus; scale bars, 5µm.

**Extended data.4.**
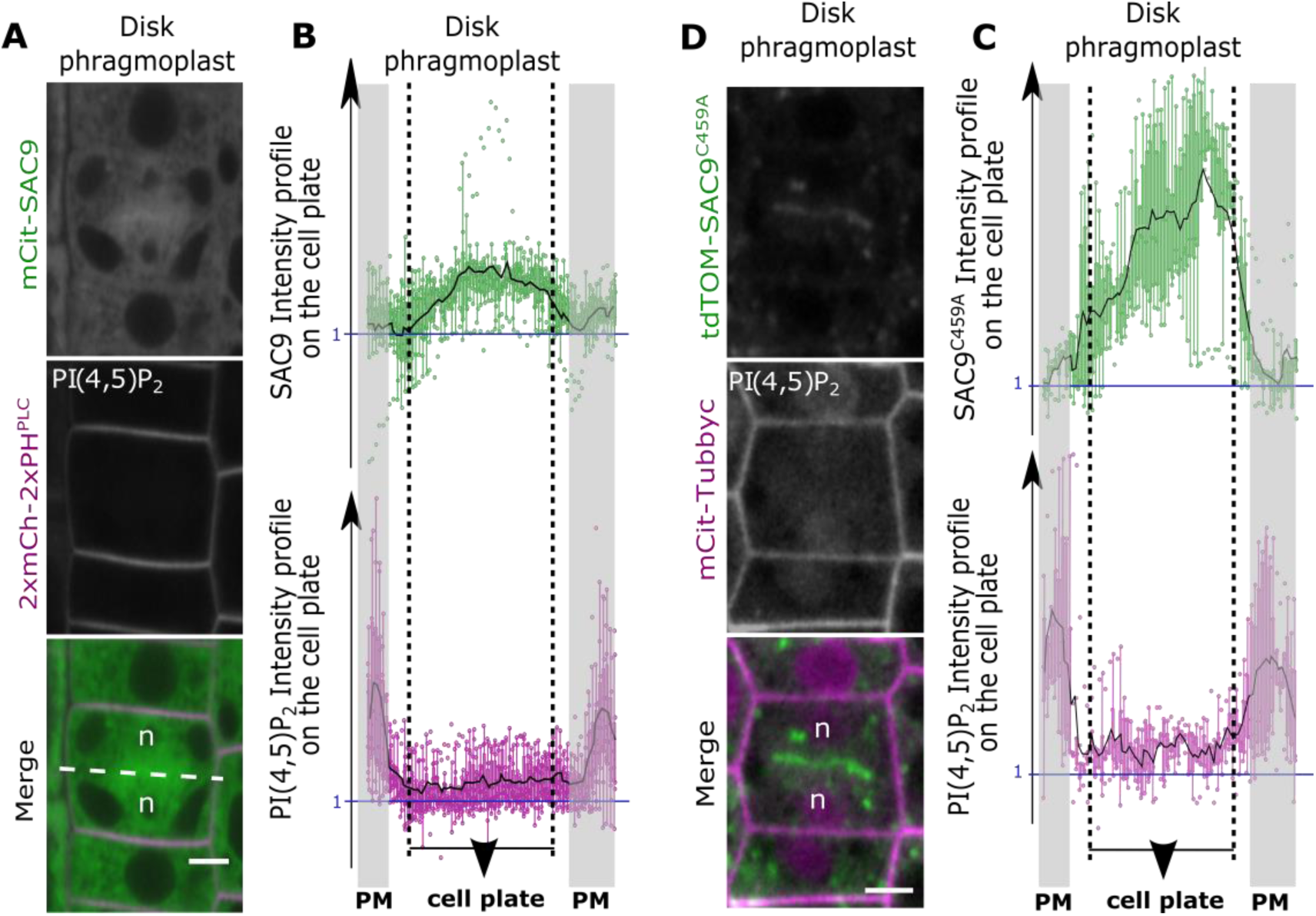
SAC9 and the PI(4,5)P_2_ localization during the disk phragmoplast stage. **a**, Confocal images of mCit-SAC9 and *2xmCh-2xPH^PLC^* at the “Disk phragmoplast” phase in WT. **b**, Normalized intensity profiles of mCit-SAC9 (top panel) and 2xmCh-2xPH^PLC^ (bottom panel) during the Disk phragmoplast phase (entire cell plate, dotted line). **c**, Confocal images of WT expressing *tdTOM-SAC9^C459A^* and *mCit-Tubbyc*. **d**, Normalized intensity profiles of tdTOM-SAC9^C459A^ (top panel) and mCit-Tubbyc (bottom panel).

**Extended data.5.**
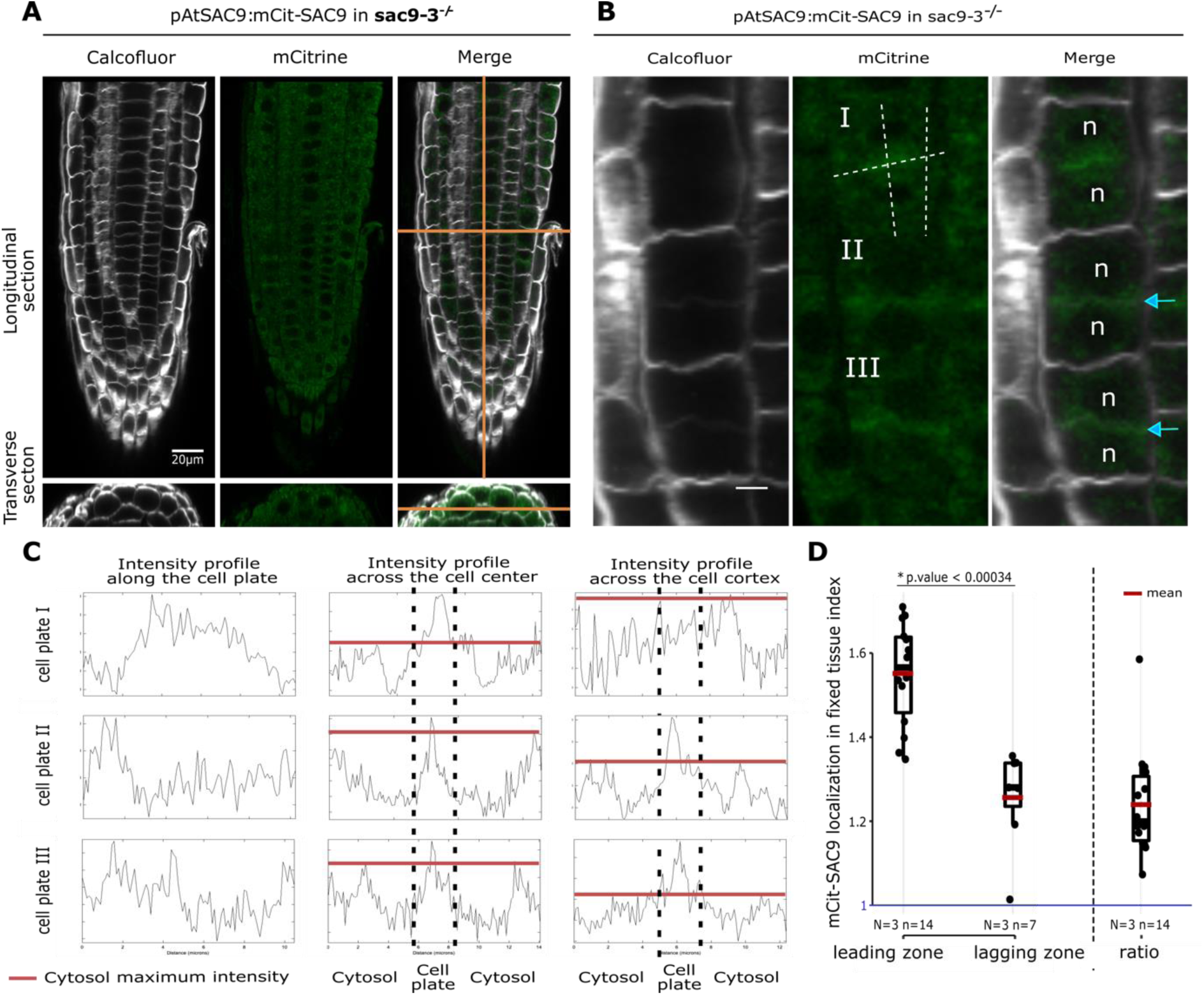
mCit-SAC9 subcellular localization and fixed tissue. **a**, Representative images of mCit-SAC9 signal obtained after fixation of the tissue and calcofluor labeling of the cell wall. Cortex cell layer is presented. (Scale bar 20µm). **b**, Example of three dividing cells of the epidermis from whole-mount fixed Arabidopsis line expressing *mCit-SAC9.* The epidermis is presented. (Scale bar 5µm). **c**, Intensity profiles of mCit-SAC9 on the cell plate axis, across the center of the cell (in the root growth axis), and finally across the cell cortex (nearby the lateral cell wall in the root growth axis). The Black dotted line delimits the cell plate, the red line corresponds to the top fluorescence intensity in the cytosol. **d**, Quantification of mCit-SAC9 localization index in fixed tissue. Blue line represents a situation in which mCit-SAC9 is not enriched compared to the cytosolic background. The cell plate accumulation edges index represents the cell plate edges fluorescence intensity divided by the cell plate center fluorescence intensity. Cell plate accumulation edges index >1 indicates a specific enrichment at the edges compared to the center of the cell plate. In the plots, middle horizontal bars represent the median, while the bottom and top of each box represent the 25^th^ and 75^th^ percentiles, respectively. At most, the whiskers extend to 1.5 times the interquartile range, excluding data beyond. For range of value under 1,5 IQR, whiskers represent the range of maximum and minimum values. Results of the statistical analysis (shown in the supplementary table) are presented (N= number of replicates, n=number of cells). Blue arrow, cell plate fusion site; n, nucleus.

**Extended data.6.**
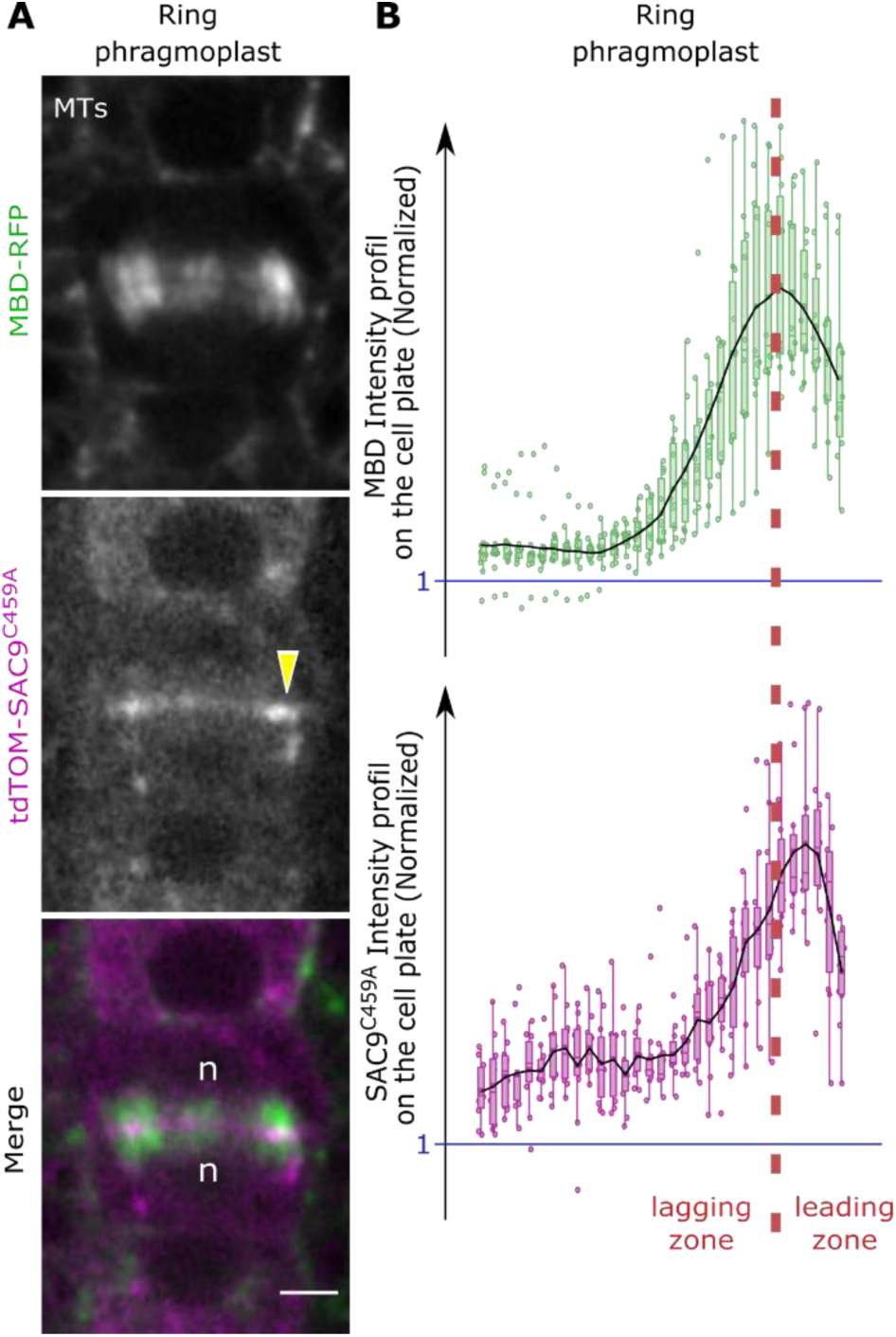
tdTOM-SAC9^C459A^ localization during the ring phragmoplast stage relative to the phragmoplast microtubules. **a**, Images of tdTOM-SAC9^C459A^ with MBD-GFP. **b**, Normalized intensity profiles corresponding to (a). Red dotted line, separation leading and lagging zone; n, nucleus; scale bar, 5µm. Yellow arrowhead, SAC9^C459A^ enrichment at the leading zone.

**Extended data.7.**
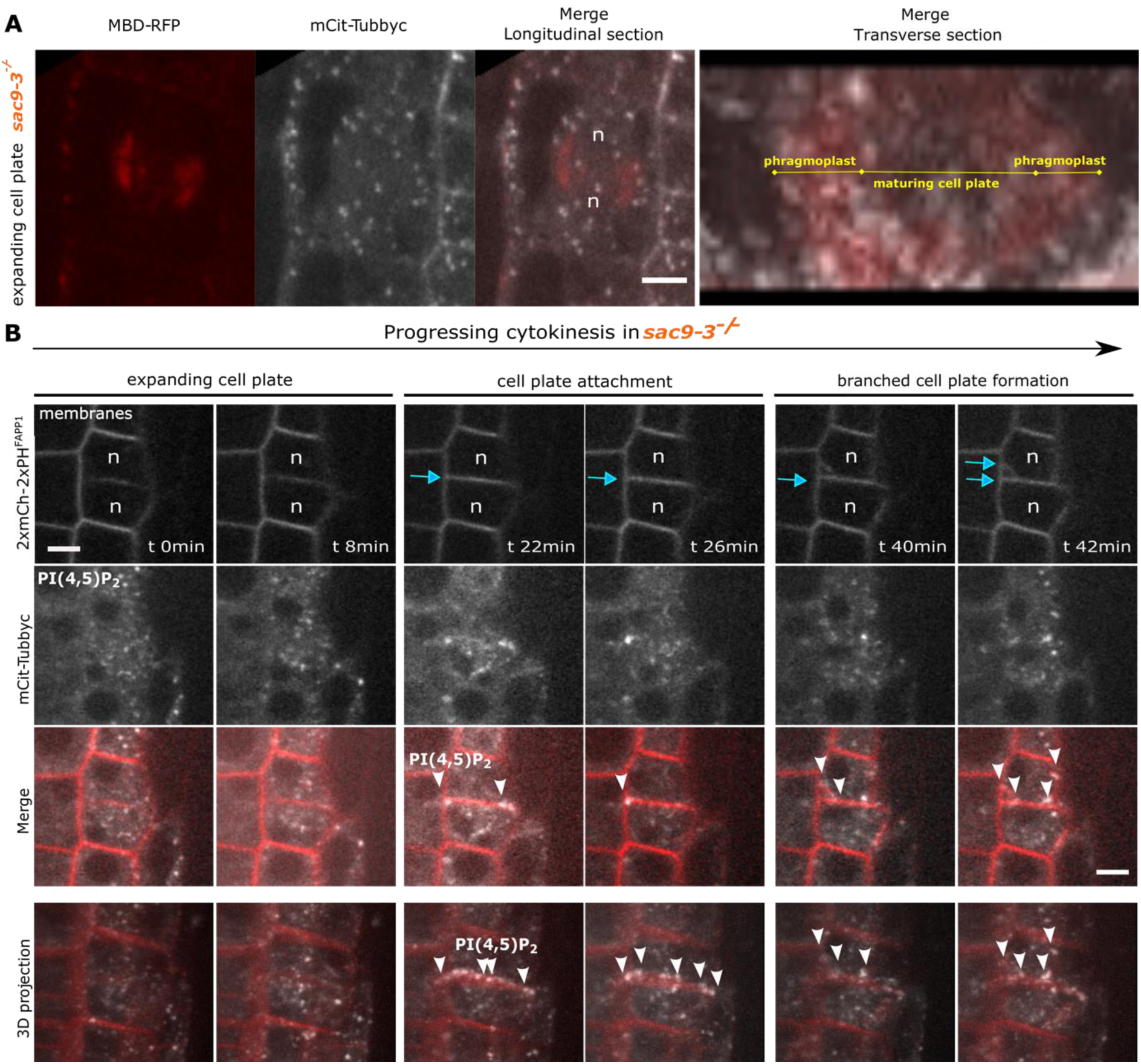
In *sac9-3*, the PI(4,5)P_2_ is abnormally recruited at the phragmoplast leading zone. **a**, Confocal images of *sac9-3* expressing *mCit-Tubbyc* and *RFP-MBD* during “the ring phragmoplast” phase or expanding cell plate phase. The first three panels correspond to a longitudinal view and the fourth panel corresponds to a transverse view of the same cell (0.7µm between optical sections). **b**, Image series and 3D projection of an *in vivo* dividing *sac9-3* cells expressing both PI(4,5)P_2_ (mCit-Tubbyc, white) and PI(4)P (mCh-PH^FAPP1^, red) biosensors. Images were taken every 2 min (between 20 and 30 Z-stack per frame) for 2 h with a root tracking system (*38*). Note that here, 2xmCh-2xPH^FAPP1^ is used at the membrane marker (plasma membrane and cell plate). Blue arrow, cell plate fusion site; n, the nucleus; white arrowhead, in sac9-3, PI(4,5)P_2_ biosensor “ring” surrounding the cell plate and PI(4,5)P_2_ biosensor spot where the branch emerged and at its tip. Scale bars, 5 µm.

**Extended data.8.**
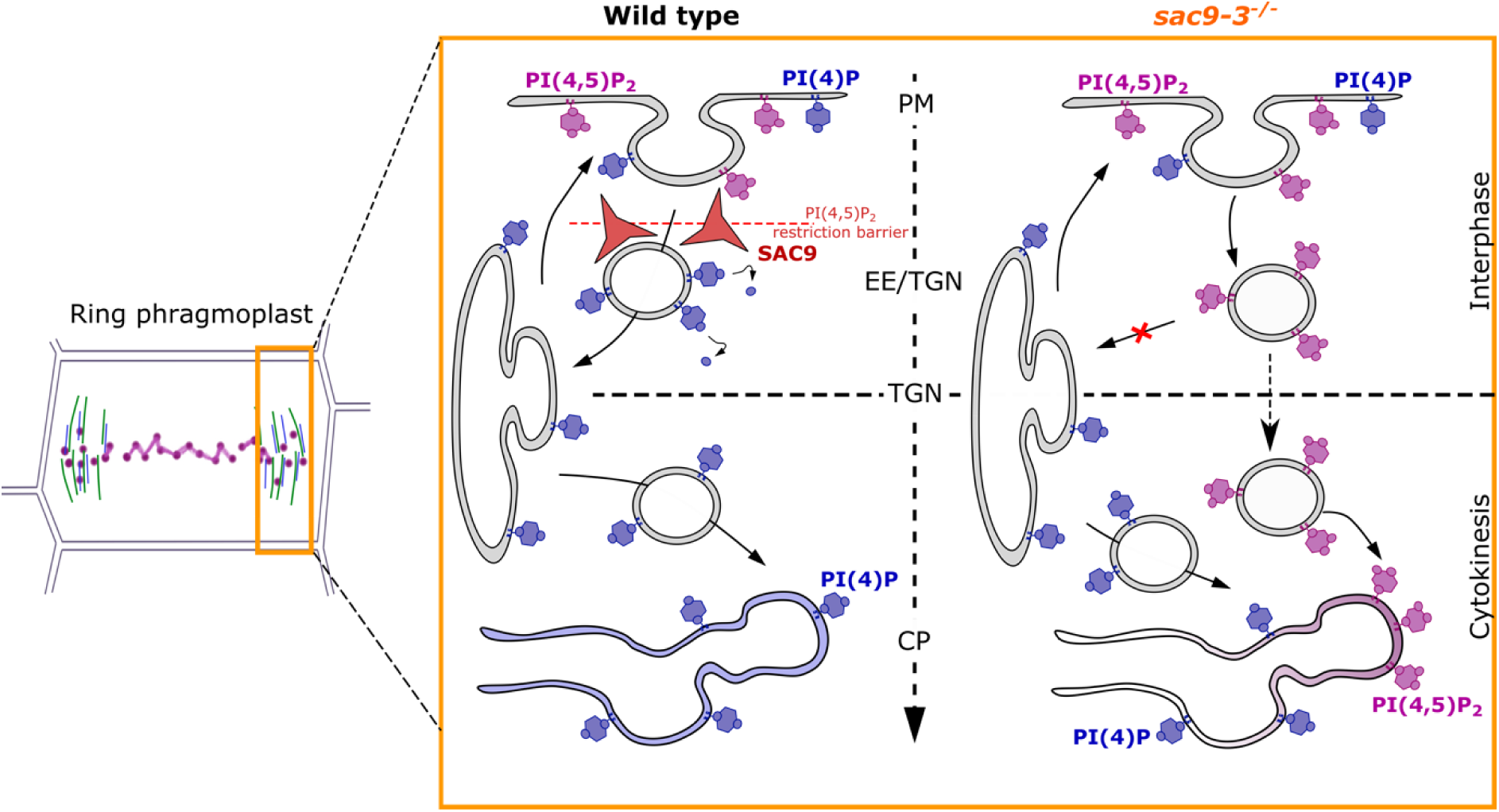
Model for the abnormal recruitment of PI(4,5)P_2_ in *sac9-3*. Hypothetical model of the mechanism at the cell plate edges based on our results. In wild-type plants, SAC9 restricts the PI(4,5)P_2_ at the plasma membrane giving rise to a endomembrane system free of this lipid. Upon SAC9 absence, the PI(4,5)P_2_ is accumulated in endomembrane compartments lost in the endocytic pathway, no longer able to reach and fuse to the TGN. This abnormal PI(4,5)P_2_ containing membranes accumulating in absence of SAC9 are during the cytokinetic process integrated at the leading edges of the phragmoplast notably during its fusion to the plasma membrane.

**Extended data.9.**
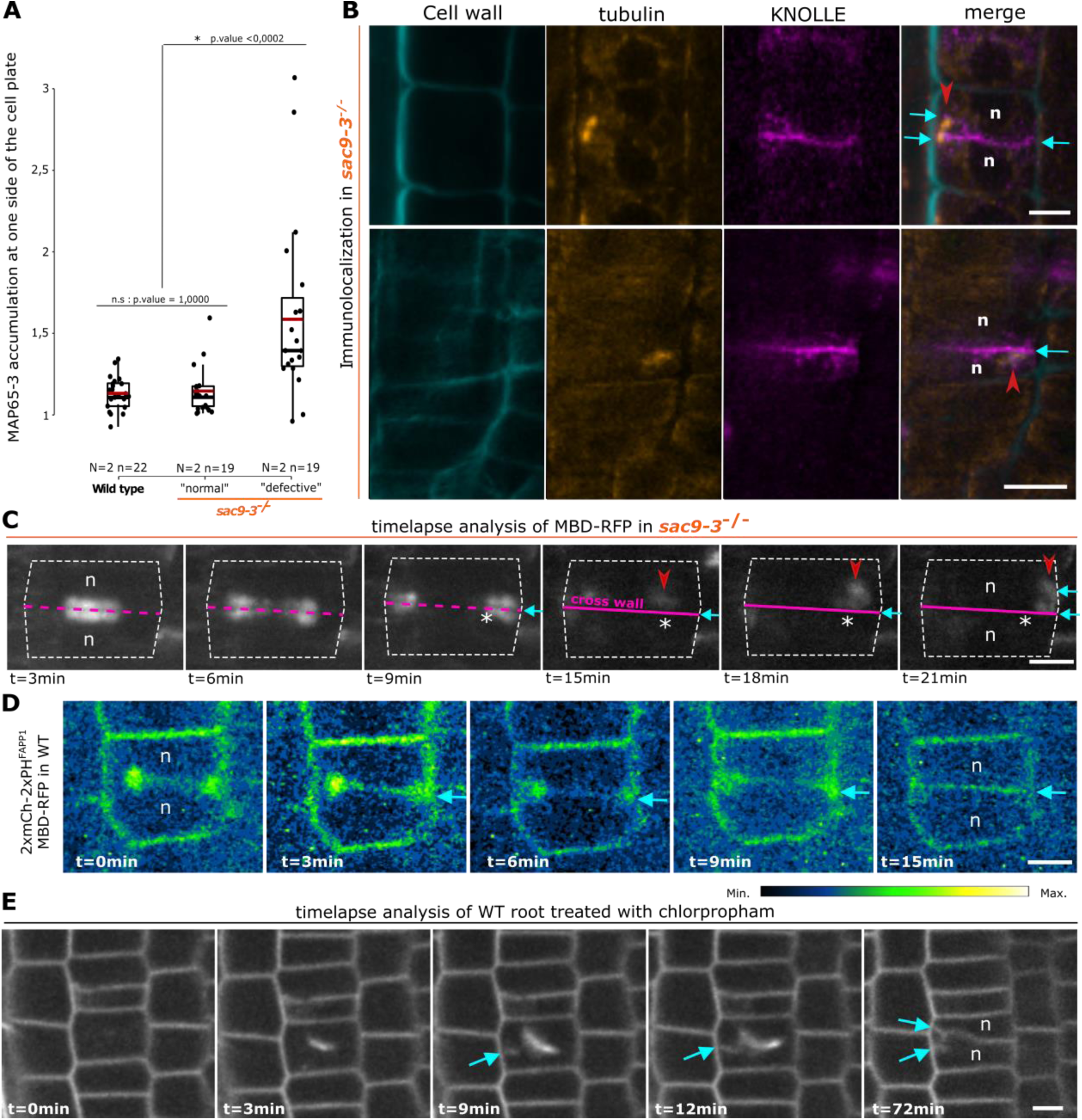
In *sac9-3*, cell plate branching correlates with microtubule perturbation. **a**, index of MAP65-3 accumulation at one side of the cell plate. In the plots, middle horizontal bars represent the median, while the bottom and top of each box represent the 25^th^ and 75^th^ percentiles, respectively. At most, the whiskers extend to 1.5 times the interquartile range, excluding data beyond. For range of value under 1,5 IQR, whiskers represent the range of maximum and minimum values. Results of the statistical analysis (shown in the supplementary table) are presented (N = number of replicates, n = number of cells). **b**, Co-immunolocalization against αtubulin and KNOLLE in *sac9-3* stained with calcofluor. Scale bar, 5 μm. **c**, Timelapse of a Z projection of RFP-MBD in *sac9-3*. Scale bar, 5 μm. **d**, Z projection of 2xmCherry-2xPH^FAPP1^ and RFP-MBD in WT. Here, the signal is color coded in green fire blue (see scale bar on the right). The signal corresponding to microtubules and the one corresponding to membrane are not distinguished). Note that here, 2xmCh-2xPH^FAPP1^ is used at the membrane marker (plasma membrane and cell plate). **e**, Image series acquired with a root tracking system of WT treated for 2 h with chlorpropham 20 µM. Blue arrow, cell plate fusion site; Red arrowhead, branched cell plate; n, nucleus; Asterik, position at the cell plate for the branch’s emergence. Scale bar, 10 μm.

**Extended data.10.**
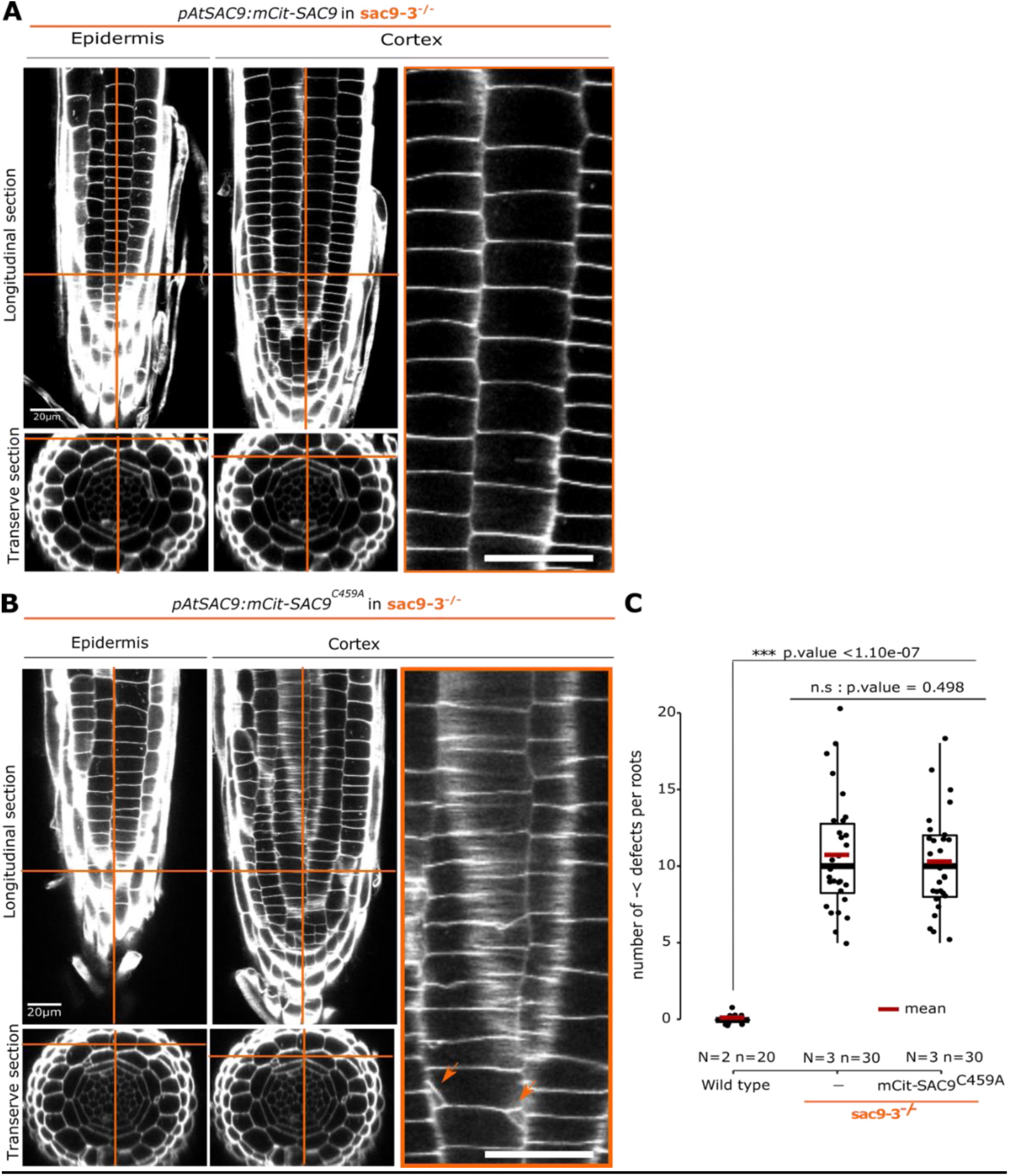
Mutation in the putative catalytic domain of SAC9 does not rescue the phenotype during cell division. **a, b**, Z-stack images of calcofluor-stained fixed roots (7-day-old) *sac9-3* expressing the functional *mCit-SAC9* (a) or *mCit-SAC9^C459A^* (b). The left panel shows the epidermis layer in a longitudinal section (top, scale bar 20 µm.) and a transverse section (bottom). The middle panel shows the cortex layer, in the same way as the epidermis. The right panel shows a crop of the cortex layer (scale bar 5 µm). Orange arrow, cell division defects; Orange cross-section, position within the tissue for longitudinal and transverse sections. **c**, Quantification of the number of cell wall defects in wild-type, *sac9-3*, and mCit-SAC9^C459A^ in *sac9-3*. In the plots, middle horizontal bars represent the median, while the bottom and top of each box represent the 25^th^ and 75^th^ percentiles, respectively. At most, the whiskers extend to 1.5 times the interquartile range, excluding data beyond. For range of value under 1,5 IQR, whiskers represent the range of maximum and minimum values. Results of the statistical analysis (shown in the supplementary table) are presented (N = number of replicates, n = number of cells).

**Extended data.11.**
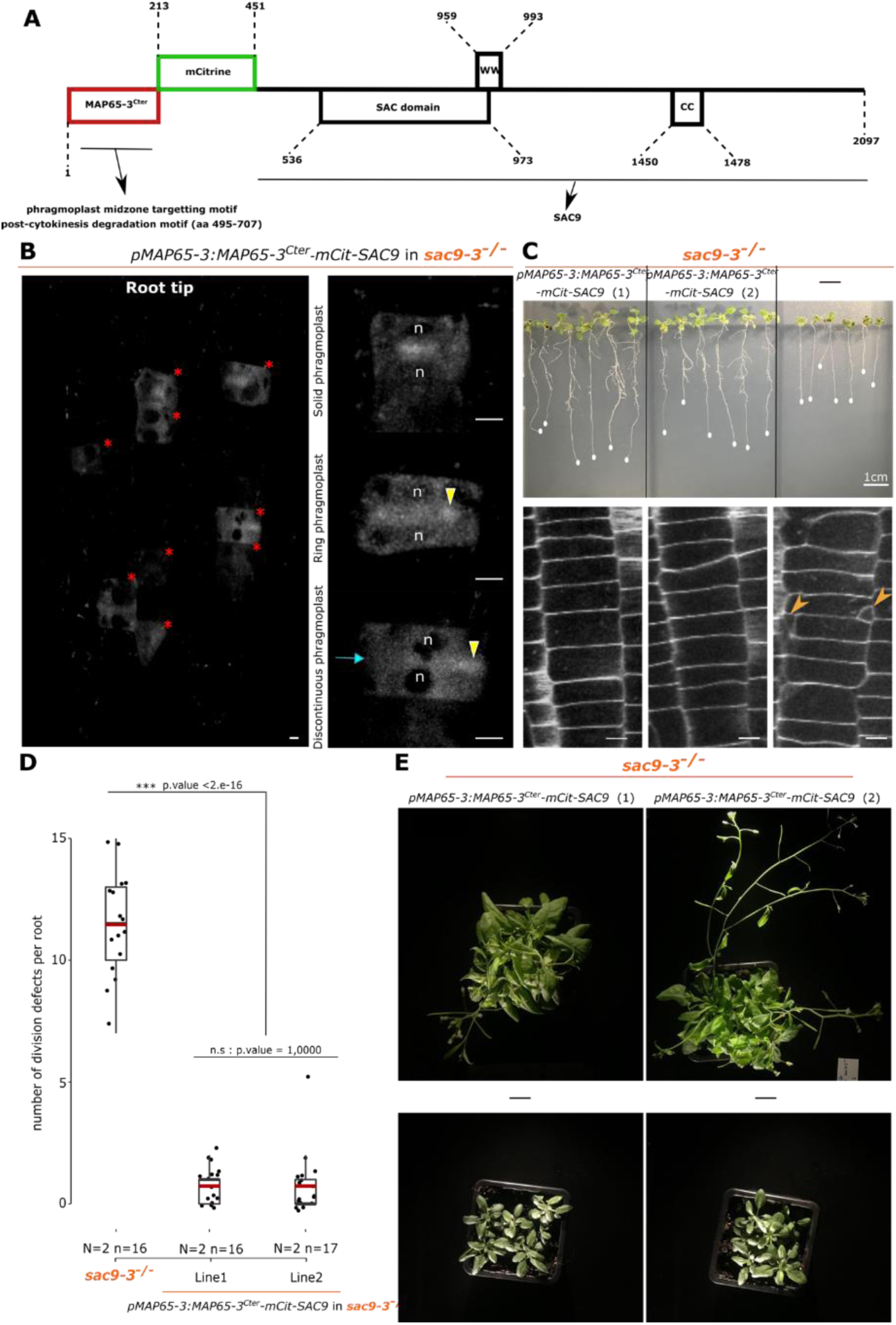
Partial rescue of *sac9-3* when SAC9 is only present in dividing cells. **a**, Diagram representing the 2097 amino-acid MAP65-3^Cter^ -mCit-SAC9 protein. We fused the C-terminal (Cter) domain of MAP65-3 that contains a microtubule binding site that enables specific activity in the phragmoplast midzone and its degradation post cytokinesis (*24*) to mCIT and SAC9 genomic sequence. **b,** left: Expression of *pMAP65-3:MAP65-3*^Cter^ *-mCit-SAC9* in *sac9-3* dividing cells (red star) of the root meristematic zone. Right: localization of MAP65-3^cter^ -mCit-SAC9 at three cytokinesis steps. Scale bars, 5µm. **c**, Phenotypes of *sac9-3* (right panel) complemented with two MAP65-3^Cter^-mCit-SAC9 independent lines (middle and left panels). Top panel: representative images of the macroscopic phenotype observed (21dpi); Bottom panel: images of calcofluor-stained fixed roots (10 day-old). Scale bars, 5µm. Orange arrowhead, division defects; n, nucleus; yellow arrowhead, enrichment on the cell plate; Blue arrow, the cell plate fusion site. **d**, Quantification of the number of division defects in *sac9-3* and *sac9-3* expressing *MAP65-3:MAP65-3^Cter^-mCit-SAC9*. In the plots, middle horizontal bars represent the median, while the bottom and top of each box represent the 25^th^ and 75^th^ percentiles, respectively. At most, the whiskers extend to 1.5 times the interquartile range, excluding data beyond. For range of value under 1,5 IQR, whiskers represent the range of maximum and minimum values. Results of the statistical analysis (shown in the supplementary table) are presented (N= number of replicates, n=number of cells). **e**, Representative images of the aerial phenotype observed (6 week old plants).

**Extended data.12.**
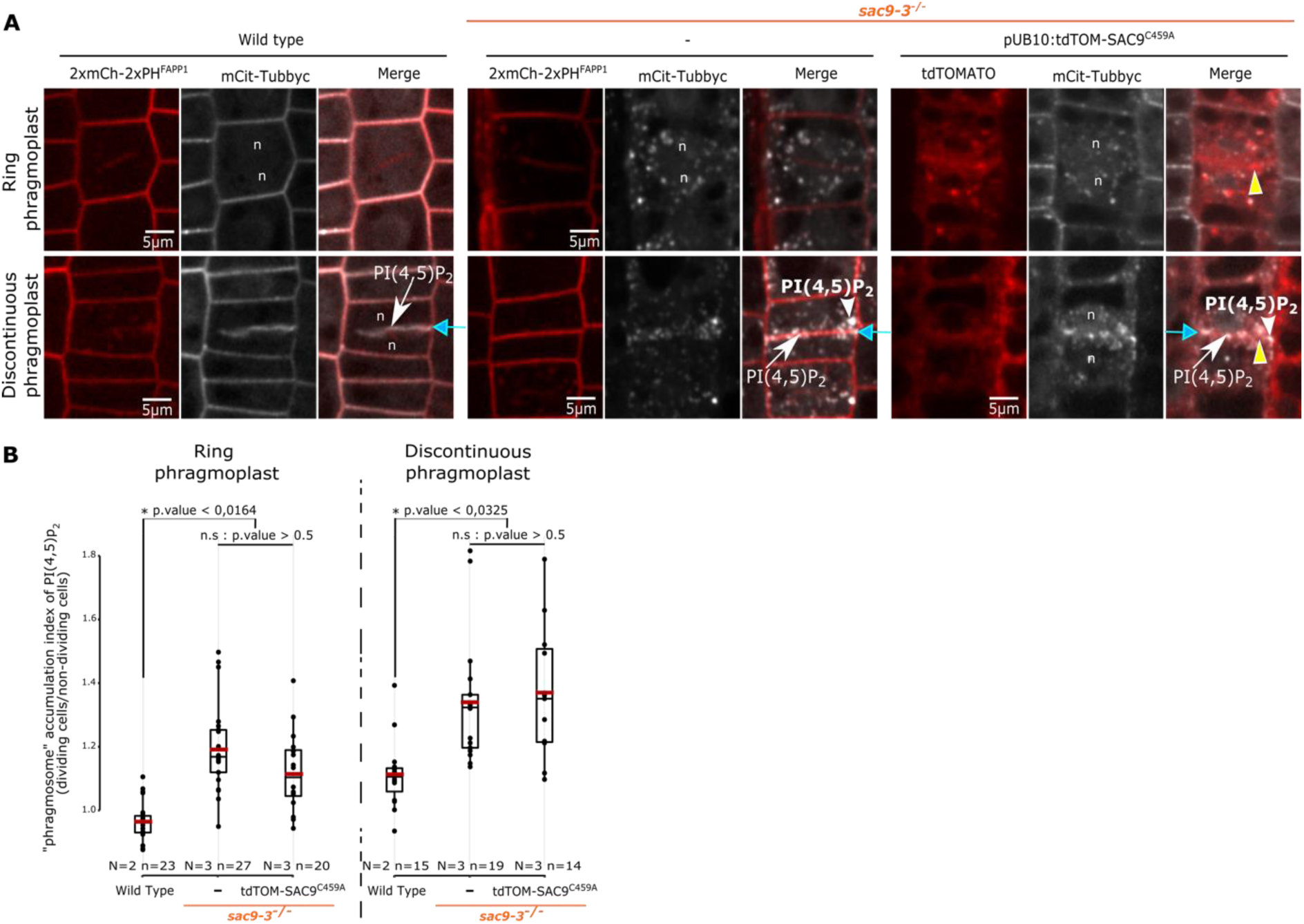
Mutation in the putative catalytic domain of SAC9 does not rescue the PI(4,5)P_2_ biosensor misregulation during cell division. **a**, Confocal images of WT expressing *2xmCh-2xPH^FAPP1^ (red)*, *sac9-3* expressing *2xmCh-2xPH^FAPP1^ (red)* and *sac9-3* expressing *tdTOM-SAC9^C459A^*, together with *mCit-Tubbyc* (white). For the sets of three panels, the top images correspond to the “expanding cell plate” phase and the bottom image to the “partially attached cell plate” phase. Here, 2xmCh-2xPH^FAPP1^ is used as a marker for membranes (plasma membrane and cell plate). The white arrow points out the cell plate for PI(4,5)P_2_, yellow arrowhead the SAC9^C459A^ enrichment and “n” the nucleus (scale bars, 5µm). b, Comparison of the phragmosome accumulation index for PI(4,5)P_2_ biosensors (mCit-Tubbyc) at the “expanding cell plate” (left panel) and partially attached cell plate” (right panel) between WT, *sac9-3* mutant and *sac9-3* expressing *tdTOM-SAC9^C459A^*. In the plots, middle horizontal bars represent the median, while the bottom and top of each box represent the 25^th^ and 75^th^ percentiles, respectively. At most, the whiskers extend to 1.5 times the interquartile range, excluding data beyond. For range of value under 1,5 IQR, whiskers represent the range of maximum and minimum values. Results of the statistical analysis (shown in the supplementary table) are presented (N = number of replicates, n = number of cells).

**Extended data.13.**
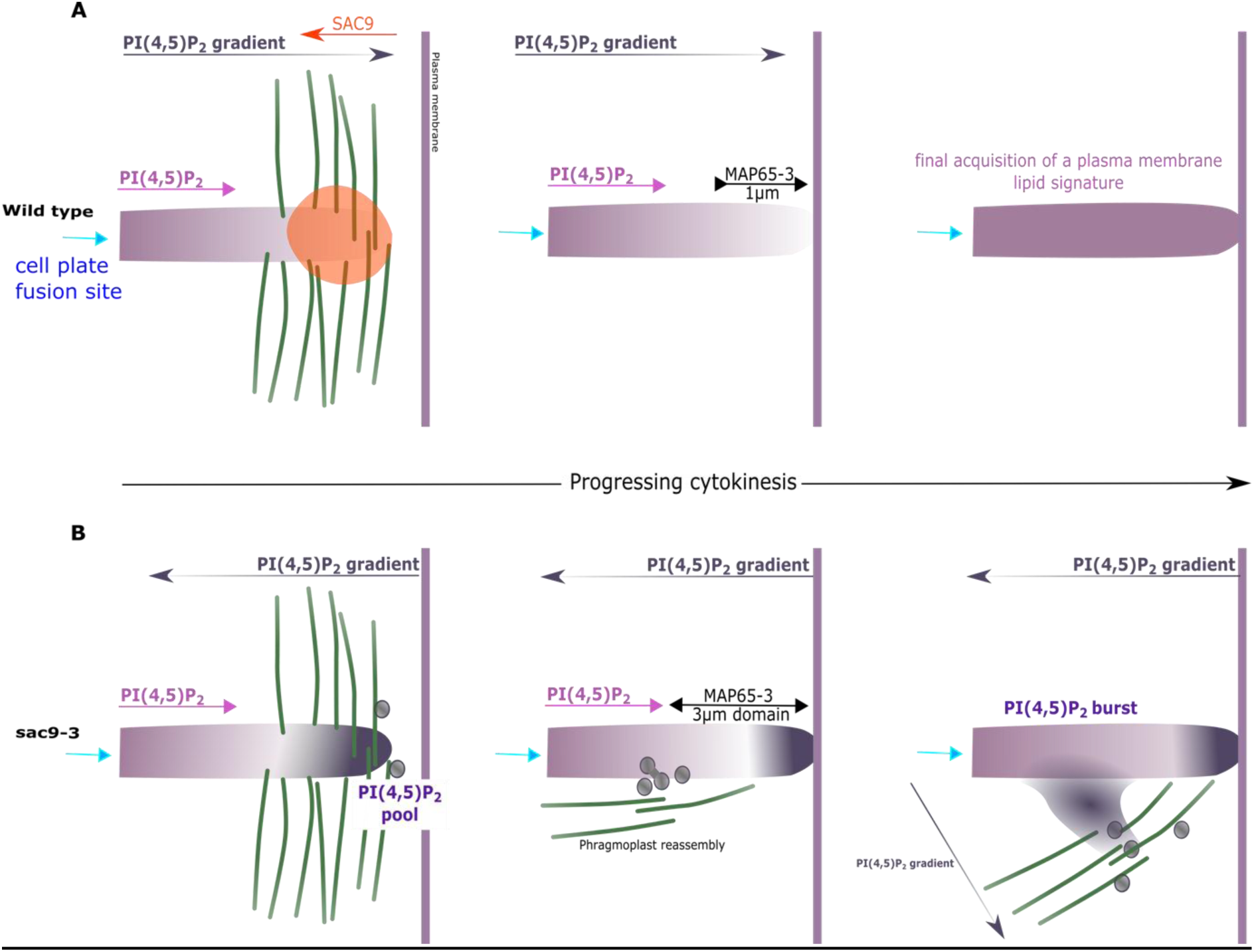
Proposed model. **a**, Upon a first partial cell plate fusion to the plasma membrane, during discontinuous phragmoplast stage, the remaining expanding phragmoplast has to accommodate rapid PI(4,5)P_2_ transfer at the cell plate. This PI(4,5)P_2_ is tightly controlled on the expanding phragmoplast, through the action of SAC9 enzyme, leading to the formation of a PI(4,5)P_2_ gradient from the fused cell plate attachment (left part of the cell plate) toward the expanding edges. Concomitantly to the phragmoplast insertion to the plasma membrane, this gradient progressively reaches the entire cell plate, accompanied by a decrease of MAP65-3 domain, and marks the final cell plate maturation and formation of a new cross wall. **b**, In *sac9* mutant, PI(4,5)P_2_ gradient on the discontinuous phragmoplast is in the opposite direction relative to the cell plate growth direction, creating a conflict in the spatial coordination of the expanding leading zone. Until the cell plate’s complete attachment to the plasma membrane, this misoriented gradient has no impact on the cell plate’s outward progression, which is driven by cortical division zone-phragmoplast leading zone proteins. Following cell plate fusion to the plasma membrane, and phragmoplast component disengage, leaving the PI(4,5)P_2_ gradient’s as the only remaining polar factor cue. Mis-polarized PI(4,5)P_2_ gradient in *sac9* is perceived by MAP65-3 as a spatial cue leading first to MAP65-3 ∼3µm domain stabilization, even after cell plate complete attachment, and second to reassembly of the phragmoplast apparatus on the inner side of the MAP65-3 domain. Following phragmoplast reassembly, a cell plate protuberance emerges and expands toward the plasma membrane followed by a PI(4,5)P_2_ burst, generating a second gradient orienting the branch expansion.

**Extended video.1** Colocalization analysis of tdTOM-SAC9^C459A^ (magenta) with MAP65-3-GFP (green)

Time-course analysis during cytokinesis of tdTOM-SAC9^C459A^ (magenta) with MAP65-3-GFP (green) using a root tracking system (*38*). The division was imaged every two minutes. Scale bar, 5µm

**Extended video.2** PI(4,5)P_2_ behavior during cytokinesis in *sac9-3* mutant

Time-course analysis using a root tracking system (*38*) of mCit-Tubbyc (white) and 2xmCH-2xPH^FAPP1^ (red) subcellular localization during cytokinesis in *sac9-3*. Here, 2xmCh-2xPH^FAPP1^ is used as a marker for membranes (plasma membrane and cell plate). The division was imaged every two minutes. Scale bar: 5 µm.

**Extended video.3** MAP65-3-GFP behavior during cytokinesis in wild-type plants

Time-course analysis using a root tracking system (*38*) of MAP65-3-GFP subcellular localization during cytokinesis in WT. The division was imaged every two minutes. Scale bar : 5µm

**Extended video.4 and 5** MAP65-3-GFP behavior’s during cytokinesis in *sac9-3*

Exemple of defective cytokinesis in *sac9-3* expressing *MAP65-3-GFP* and imaged over time using a root tracking system (*38*). The division was imaged every two minutes. scale bar: 5 µm

**Extended video.6** Mild treatment with a microtubule perturbator phenocopies *sac9-3* phenotype

Time-course analysis using a root tracking system of the plasma membrane labeling PI4P biosensor (mCIT-P4M) in WT root treated for 2h with chlorpropham. Here, mCIT-P4M is used as a marker for membranes (plasma membrane and cell plate). Scale bar: 10 µm

**Extended video.7** Imaging of the defects during cytokinesis in *sac9-3*

Time-course analysis using a root tracking system (*38*) of 2xmCH-2xPH^FAPP1^ in *sac9-3* during cell plate defect formation (arrow). Here, 2xmCH-2xPH^FAPP1^ is used as a marker for membranes (plasma membrane and cell plate). Scale bar: 5 µm

**Extended video.8** branched cell wall defects segmentation in *sac9-3*

Rotation of a branched cell wall defect (green) and its neighboring cell (gray) 3D view imaged after calcofluor staining.

**Extended video.9** branched cell wall defects segmentation in *sac9-3*

Rotation of branched cell wall defects accumulation (green) and their neighboring cell (gray) 3D view imaged after calcofluor staining.

**Extended video.10** PI(4,5)P_2_ behavior during double cell plate attachment in *sac9-3*

Time-course analysis using a root tracking system (*38*) of mCit-Tubbyc (white) and 2xmCH-2xPH^FAPP1^ (red) subcellular localization during cytokinesis in *sac9-3*. Here, 2xmCH-2xPH^FAPP1^ is used as a marker for membranes (plasma membrane and cell plate). The division was imaged every two minutes. Scale bar, 5µm

**Supplementary tables** Resources and statistics.

**a**, Reagent and resources. **b,** Details of the statistics corresponding to Figure 1f. **c,** Details of the statistics corresponding to Figure 2c, 2f. **d,** Details of the statistics corresponding to Figure 3d, 3f. **e,** Details of the statistics corresponding to Figure 5b. **f,** Details of the statistics corresponding to Figure 6b. **g,** Details of the statistics corresponding to Extended data 3b, 3d. **h,** Details of the statistics corresponding to Extended data 5d. **i,** Details of the statistics corresponding to Extended data 9a. **j,** Details of the statistics corresponding to Extended data 10c. **k,** Details of the statistics corresponding to Extended data 11d **l,** Details of the statistics corresponding to Extended data 12b.

## REFERENCES

1. A. Smertenko, F. Assaad, F. Baluska, M. Bezanilla, H. Buschmann, G. Drakakaki, M. T. Hauser, M. Janson, Y. Mineyuki, I. Moore, S. Muller, T. Murata, M. S. Otegui, E. Panteris, C. Rasmussen, A. C. Schmit, J. Samaj, L. Samuels, L. A. Staehelin, D. Van Damme, G. Wasteneys, V. Zarsky, Plant Cytokinesis: Terminology for Structures and Processes. Trends Cell Biol. 27, 885–894 (2017).

2. P. Livanos, S. Muller, Division Plane Establishment and Cytokinesis. Annual review of plant biology. 70, 239–267 (2019).

3. J. Chang-Jie, S. Sonobe, Identification and preliminary characterization of a 65 kDa higher-plant microtubule-associated protein. Journal of Cell Science. 105, 891–901 (1993).

4. Y. R. Lee, B. Liu, Identification of a phragmoplast-associated kinesin-related protein in higher plants. Current biology : CB. 10, 797–800 (2000).

5. P. J. Moore, L. A. Staehelin, Immunogold localization of the cell-wall-matrix polysaccharides rhamnogalacturonan I and xyloglucan during cell expansion and cytokinesis inTrifolium pratense L.; implication for secretory pathways. Planta. 174, 433–445 (1988).

6. A. L. Samuels, T. H. Giddings, L. A. Staehelin, Cytokinesis in tobacco BY-2 and root tip cells: a new model of cell plate formation in higher plants. J Cell Biol. 130, 1345–57 (1995).

7. J. M. Segui-Simarro, J. R. Austin, E. A. White, L. A. Staehelin, Electron tomographic analysis of somatic cell plate formation in meristematic cells of Arabidopsis preserved by high-pressure freezing. The Plant cell. 16, 836–56 (2004).

8. K. Rybak, A. Steiner, L. Synek, S. Klaeger, I. Kulich, E. Facher, G. Wanner, B. Kuster, V. Zarsky, S. Persson, F. F. Assaad, Plant cytokinesis is orchestrated by the sequential action of the TRAPPII and exocyst tethering complexes. Dev Cell. 29, 607–20 (2014).

9. L. C. Noack, Y. Jaillais, Functions of Anionic Lipids in Plants. Annual review of plant biology. 71, 71– 102 (2020).

10. M. Heilmann, I. Heilmann, Regulators regulated: Different layers of control for plasma membrane phosphoinositides in plants. Current Opinion in Plant Biology. 67, 102218 (2022).

11. A. Lebecq, M. Doumane, A. Fangain, V. Bayle, J. X. Leong, F. Rozier, M. del Marques-Bueno, L. Armengot, R. Boisseau, M. L. Simon, The Arabidopsis SAC9 enzyme is enriched in a cortical population of early endosomes and restricts PI (4, 5) P2 at the plasma membrane. Elife. 11, e73837 (2022).

12. P. Marhava, A. C. A. Fandino, S. W. Koh, A. Jelínková, M. Kolb, D. P. Janacek, A. S. Breda, P. Cattaneo, U. Z. Hammes, J. Petrášek, Plasma membrane domain patterning and self-reinforcing polarity in Arabidopsis. Developmental cell. 52, 223–235 (2020).

13. Y. Mei, W.-J. Jia, Y.-J. Chu, H.-W. Xue, Arabidopsis phosphatidylinositol monophosphate 5-kinase 2 is involved in root gravitropism through regulation of polar auxin transport by affecting the cycling of PIN proteins. Cell research. 22, 581–597 (2012).

14. P. Scholz, J. Anstatt, H. E. Krawczyk, T. Ischebeck, Signalling pinpointed to the tip: the complex regulatory network that allows pollen tube growth. Plants. 9, 1098 (2020).

15. M. Doumane, A. Lebecq, L. Colin, A. Fangain, F. D. Stevens, J. Bareille, O. Hamant, Y. Belkhadir, T. Munnik, Y. Jaillais, M.-C. Caillaud, Inducible depletion of PI(4,5)P2 by the synthetic iDePP system in Arabidopsis. Nature Plants. 7, 587–597 (2021).

16. M. Fratini, P. Krishnamoorthy, I. Stenzel, M. Riechmann, M. Matzner, K. Bacia, M. Heilmann, I. Heilmann, Plasma membrane nano-organization specifies phosphoinositide effects on Rho-GTPases and actin dynamics in tobacco pollen tubes. The Plant Cell. 33, 642–670 (2021).

17. F. Lin, P. Krishnamoorthy, V. Schubert, G. Hause, M. Heilmann, I. Heilmann, A dual role for cell plate-associated PI4Kbeta in endocytosis and phragmoplast dynamics during plant somatic cytokinesis. The EMBO journal. 38 (2019), doi:10.15252/embj.2018100303.

18. M. L. Simon, M. P. Platre, M. M. Marques-Bueno, L. Armengot, T. Stanislas, V. Bayle, M. C. Caillaud, Y. Jaillais, A PtdIns(4)P-driven electrostatic field controls cell membrane identity and signalling in plants. Nat Plants. 2, 16089 (2016).

19. W. Van Leeuwen, J. E. Vermeer, T. W. Gadella Jr, T. Munnik, Visualization of phosphatidylinositol 4, 5-bisphosphate in the plasma membrane of suspension-cultured tobacco BY-2 cells and whole Arabidopsis seedlings. The Plant Journal. 52, 1014–1026 (2007).

20. M. L. Simon, M. P. Platre, S. Assil, R. van Wijk, W. Y. Chen, J. Chory, M. Dreux, T. Munnik, Y. Jaillais, A multi-colour/multi-affinity marker set to visualize phosphoinositide dynamics in Arabidopsis. Plant J. 77, 322–37 (2014).

21. M. Doumane, A. Lebecq, A. Fangain, V. Bayle, F. Rozier, M. del M. Marquès-Bueno, R. P. Boisseau, M. L. A. Simon, L. Armengot, Y. Jaillais, M.-C. Caillaud, bioRxiv, in press, doi:10.1101/2021.09.10.459735.

22. K. L. Farquharson, MAP65-3 cross-links interdigitated microtubules in the phragmoplast. The Plant cell. 23, 2807 (2011).

23. C. M. Ho, T. Hotta, F. Guo, R. W. Roberson, Y. R. Lee, B. Liu, Interaction of antiparallel microtubules in the phragmoplast is mediated by the microtubule-associated protein MAP65-3 in Arabidopsis. The Plant cell. 23, 2909–23 (2011).

24. C. M. Ho, Y. R. Lee, L. D. Kiyama, S. P. Dinesh-Kumar, B. Liu, Arabidopsis microtubule-associated protein MAP65-3 cross-links antiparallel microtubules toward their plus ends in the phragmoplast via its distinct C-terminal microtubule binding domain. The Plant cell. 24, 2071–85 (2012).

25. M. Sasabe, K. Kosetsu, M. Hidaka, A. Murase, Y. Machida, Arabidopsis thaliana MAP65-1 and MAP65-2 function redundantly with MAP65-3/PLEIADE in cytokinesis downstream of MPK4. Plant signaling & behavior. 6, 743–7 (2011).

26. M.-C. Caillaud, Tools for studying the cytoskeleton during plant cell division. Trends in Plant Science (2022).

27. Y. Boutte, M. Frescatada-Rosa, S. Men, C. M. Chow, K. Ebine, A. Gustavsson, L. Johansson, T. Ueda, I. Moore, G. Jurgens, M. Grebe, Endocytosis restricts Arabidopsis KNOLLE syntaxin to the cell division plane during late cytokinesis. The EMBO journal. 29, 546–58 (2010).

28. E. P. Eleftheriou, T. I. Baskin, P. K. Hepler, Aberrant cell plate formation in the Arabidopsis thaliana microtubule organization 1 mutant. Plant & cell physiology. 46, 671–5 (2005).

29. E. Eleftheriou, E. Bekiari, Ultrastructural effects of the herbicide chlorpropham (CIPC) in root tip cells of wheat. Plant and soil. 226, 11–19 (2000).

30. D. Dambournet, M. Machicoane, L. Chesneau, M. Sachse, M. Rocancourt, A. El Marjou, E. Formstecher, R. Salomon, B. Goud, A. Echard, Rab35 GTPase and OCRL phosphatase remodel lipids and F-actin for successful cytokinesis. Nature cell biology. 13, 981–8 (2011).

31. C. Cauvin, M. Rosendale, N. Gupta-Rossi, M. Rocancourt, P. Larraufie, R. Salomon, D. Perrais, A. Echard, Rab35 GTPase Triggers Switch-like Recruitment of the Lowe Syndrome Lipid Phosphatase OCRL on Newborn Endosomes. Current biology : CB. 26, 120–8 (2016).

32. I. Kouranti, M. Sachse, N. Arouche, B. Goud, A. Echard, Rab35 regulates an endocytic recycling pathway essential for the terminal steps of cytokinesis. Current biology : CB. 16, 1719–25 (2006).

33. M. C. Caillaud, P. Lecomte, F. Jammes, M. Quentin, S. Pagnotta, E. Andrio, J. de Almeida Engler, N. Marfaing, P. Gounon, P. Abad, B. Favery, MAP65-3 microtubule-associated protein is essential for nematode-induced giant cell ontogenesis in Arabidopsis. The Plant cell. 20, 423–37 (2008).

34. M. Karimi, A. Depicker, P. Hilson, Recombinational cloning with plant gateway vectors. Plant physiology. 145, 1144–54 (2007).

35. N. C. Shaner, R. E. Campbell, P. A. Steinbach, B. N. Giepmans, A. E. Palmer, R. Y. Tsien, Improved monomeric red, orange and yellow fluorescent proteins derived from Discosoma sp. red fluorescent protein. Nature biotechnology. 22, 1567–1572 (2004).

36. C. M. Ho, Y. R. Lee, L. D. Kiyama, S. P. Dinesh-Kumar, B. Liu, Arabidopsis microtubule-associated protein MAP65-3 cross-links antiparallel microtubules toward their plus ends in the phragmoplast via its distinct C-terminal microtubule binding domain. The Plant cell. 24, 2071–85 (2012).

37. S. J. Clough, A. F. Bent, Floral dip: a simplified method for Agrobacterium-mediated transformation of Arabidopsis thaliana. The Plant Journal. 16, 735–743 (1998).

38. M. Doumane, C. Lionnet, V. Bayle, Y. Jaillais, M. C. Caillaud, Automated Tracking of Root for Confocal Time-lapse Imaging of Cellular Processes. Bio Protoc. 7 (2017), doi:10.21769/BioProtoc.2245.

39. K. Belcram, D. Legland, M. Pastuglia, “Quantification of Cell Division Angles in the Arabidopsis Root” in Plant Cell Division (Springer, 2022), pp. 209–221.

40. M. Romeiro Motta, X. Zhao, M. Pastuglia, K. Belcram, F. Roodbarkelari, M. Komaki, H. Harashima, S. Komaki, M. Kumar, P. Bulankova, B1-type cyclins control microtubule organization during cell division in Arabidopsis. EMBO reports. 23, e53995 (2022).

41. M. H. Lauber, I. Waizenegger, T. Steinmann, H. Schwarz, U. Mayer, I. Hwang, W. Lukowitz, G. Jurgens, The Arabidopsis KNOLLE protein is a cytokinesis-specific syntaxin. J Cell Biol. 139, 1485–93 (1997).

42. D. Legland, I. Arganda-Carreras, P. Andrey, MorphoLibJ: integrated library and plugins for mathematical morphology with ImageJ. Bioinformatics. 32, 3532–3534 (2016).

43. H. Wickham, ggplot2: elegant graphics for data analysis (springer, 2016).

44. D. Bates, M. Mächler, B. Bolker, S. Walker, Fitting linear mixed-effects models using lme4. arXiv preprint arXiv:1406.5823 (2014).

45. J. Fox, S. Weisberg, The car package contains functions and data sets associated with the book an R companion to applied regression. R-Project (2011).

46. T. Hothorn, F. Bretz, P. Westfall, R. M. Heiberger, R package version, in press.

47. R. Lenth, M. R. Lenth, Package ‘lsmeans.’ The American Statistician. 34, 216–221 (2018).

